# Sortilin C-terminal fragment deposition depicts tangle-related nonamyloid neuritic plaque growth in Alzheimer’s disease

**DOI:** 10.1101/2024.11.11.622955

**Authors:** Qi-Lei Zhang, Yan Wang, Sidiki Coulibaly, Zhong-Ping Sun, Xiao-Lu Cai, Tian Tu, Yu Liu, Aihua Pan, Meng-Chao Cui, Jim Manavis, Jian Wang, Yang Zhang, Xiao-Pin Wang, Xiao-Xin Yan

## Abstract

Sortilin C-terminal fragments (sorfra) can co-deposit in β-amyloid (Aβ) plaques in human brain. However, sorfra plaques develop in the cerebrum with a spatiotemporal trajectory as of tauopathy. Here we examined sorfra pathogenesis relative to neuritic plaque evolution in the human brains with amyloid and tau pathologies converged in the neocortex and hippocampus. Sorfra plaques occurred in correlation with pTau/tangle, but not Aβ, pathologies across cerebral regions, neighboring cortical/hippocampal areas, and along the sulcal valley to gyral hilltop transition. Sorfra plaques and neuritic plaques were matchable in location, shape and size between consecutive sections, and were colocalized in double-labeling preparations. Microscopical study and tissue clearance three-dimensional imaging revealed sorfra/Aβ colocalized as well as independent plaques. Among the former, sorfra labeling correlated negatively to Aβ/amyloid labeling and β-secretase-1 labeling in dystrophic neurites. Sorfra plaques were depleted of microtubule-associated protein 2 (MAP2) labeled neuronal somata and dendrites, whereas normal looking MAP2/sortilin co-labeled profiles occurred nearby. Sorfra deposits were seen in astrocytes but not microglia around the plaques. Taken together, sorfra plaques are anatomically matchable to silver stained neuritic plaques. They develop with tangle-related somatodendritic degeneration, presenting as nonamyloid growth of the Aβ plaques and formation of Aβ-independent neuritic plaques during Alzheimer’s disease pathogenesis.

## INTRODUCTION

Neuritic plaques and neurofibrillary tangles (NFTs) were discovered in human brain with silver stains over a century ago and established as the primary neuropathologies of Alzheimer’s disease (AD) during the following decades (Fiala, 2007; Walker, 2020; Beach, 2022; Iqbal, 2024). Neuritic plaques are localized lesions consisting of extracellularly deposited amorphous material and swollen neuritic processes or dystrophic neurites, often associated with reactive glial cells. Tangles appear in the forms of argyrophilic deformed somata, dystrophic neurites and neuropil threads (DeTure & Dickson, 2019). In the 1980s, β-amyloid (Aβ) peptides were identified from the cores of amyloid plaques (Glenner & Wong, 1984; Masters & Beyreuther, 1986), with phosphorylated tau (pTau) characterized as the major constitute of tangle filaments(Iqbal et al., 1986; Wolozin et al, 1986). With the application of immunohistochemistry, the dystrophic neurites were found to be axonal and dendritic in origin over the years, and enriched of the Aβ producing machinery and pTau, implicating their intrinsic link to neuritic plaque formation (Brion et al., 1991; Cai et al., 2010; Ferrer, 2023; Jordà-Siquier et al., 2022; Munoz & Wang, 1992; Sadleir et al., 2016; Su et al., 1996; Su et al., 1997; Thal et al., 1998; Wang & Munoz, 1995; Zhang et al., 2009).

Many pathogenic issues regarding Aβ and pTau accumulation relative to plaque and tangle formation are still not coherently explained (Ferrari & Sorbi, 2021; Kepp et al., 2023; Villain & Michalon, 2024). There exists a mismatch between amyloid plaques and neuritic plaques (Braak et al., 1989; Tsering & Prokop, 2024). Tangle development involves pTau accumulation but also complex molecular interplay (Duan et al., 2024; Hondius et al., 2021; Jiang et al., 2022; Moloney, Lowe, & Murray, 2021; Zhang et al., 2024). Further, it remains largely unclear as to how Aβ and pTau interact or integrate during neuritic plaque evolution (Korczyn & Grinberg, 2024; Tsering & Prokop, 2024). Currently, Aβ and pTau immunolabeling as well as silver stains are required for neuropathological diagnosis of AD (Hyman et al., 2012; Montine et al., 2012). Aβ and tau pathologies develop in the human brain following “paradoxical” spatiotemporal trajectories. Aβ deposition appears first in the isocortex (i.e., Thal phase 1), then spreads into the allocortical/limbic structures (phase 2) and further into the subcortical regions (phases 3-5) (Thal et al, 2002). Tauopathy occurs progressively in the entorhinal (Braak stages I and II), limbic (III and IV) and isocortical (V and VI) regions (Braak & Braak, 1991; Braak & Del Tredici, 2014; van der Kant et al, 2020). The convergence of the two lesions in the limbic and neocortical regions appears to be a key factor to cause neurodegeneration and dementia (Arriagada et al., 1992; Mudher & Lovestone, 2002; Serrano et al., 2016; Tiraboschi et al., 2004; Trojanowski & Lee, 2002).

We reported earlier that *sor*tilin C-terminal (CT) *fra*gments (shorted as *sorfra*) can deposit in aged and AD human brains along with Aβ as typical compact-like extracellular plaques (Hu et al., 2017; Zhou et al., 2018). However, the development of sorfra plaques in the human cerebrum progresses with a spatiotemporal pattern similar to that of tauopathy (Jiang et al., 2022; Tu et al., 2020). In other words, sorfra plaque formation exhibits some “mixed” properties of the Aβ and tau pathogenesis and appears to be an event involved in neuritic plaque evolution. Therefore, we set to address this possibility using postmortem human brains with Thal phases 3-5 Aβ and Braak stages 4-6 tau pathologies, by applying multiple combinations of co-labeling (immunolabeling, silver stain and Aβ/tau tracer labeling) along with correlative image analysis using consecutive section preparation and tissue clearance technology.

## MATERIALS AND METHODS

### Human brain samples and tissue preparation

The current study was approved by the Ethics Committee of Central South University Xiangya School of Medicine, in compliance with the Code of Ethics of the World Medical Association (Declaration of Helsinki). Brains were banked through a willed body donation program (Yan et al., 2015), with donor’s clinical records obtained whenever available. The brains were routinely assessed for neuropathological changes according to the Standard Brain Banking Protocol set by China Brain Bank Consortium(Qiu et al., 2019). A total of 22 brain samples were selected for use in the current study, with the case information, neuropathological staging and tissue usage detailed in Table 1. In brief, cryostat (35 μm) and paraffin (3-4 μm) sections from the formalin-fixed half brains were used for comparative histopathological characterizations. A formalin-fixed temporal neocortical block was obtained from an AD brain, which underwent a tissue clearance procedure, followed by fluorescent immunolabeling for Aβ and sorfra and three-dimensional (3D) imaging at low and high resolutions.

**Table 1:**
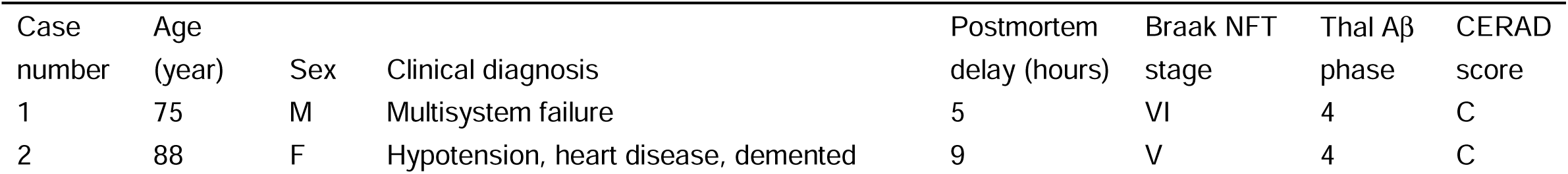
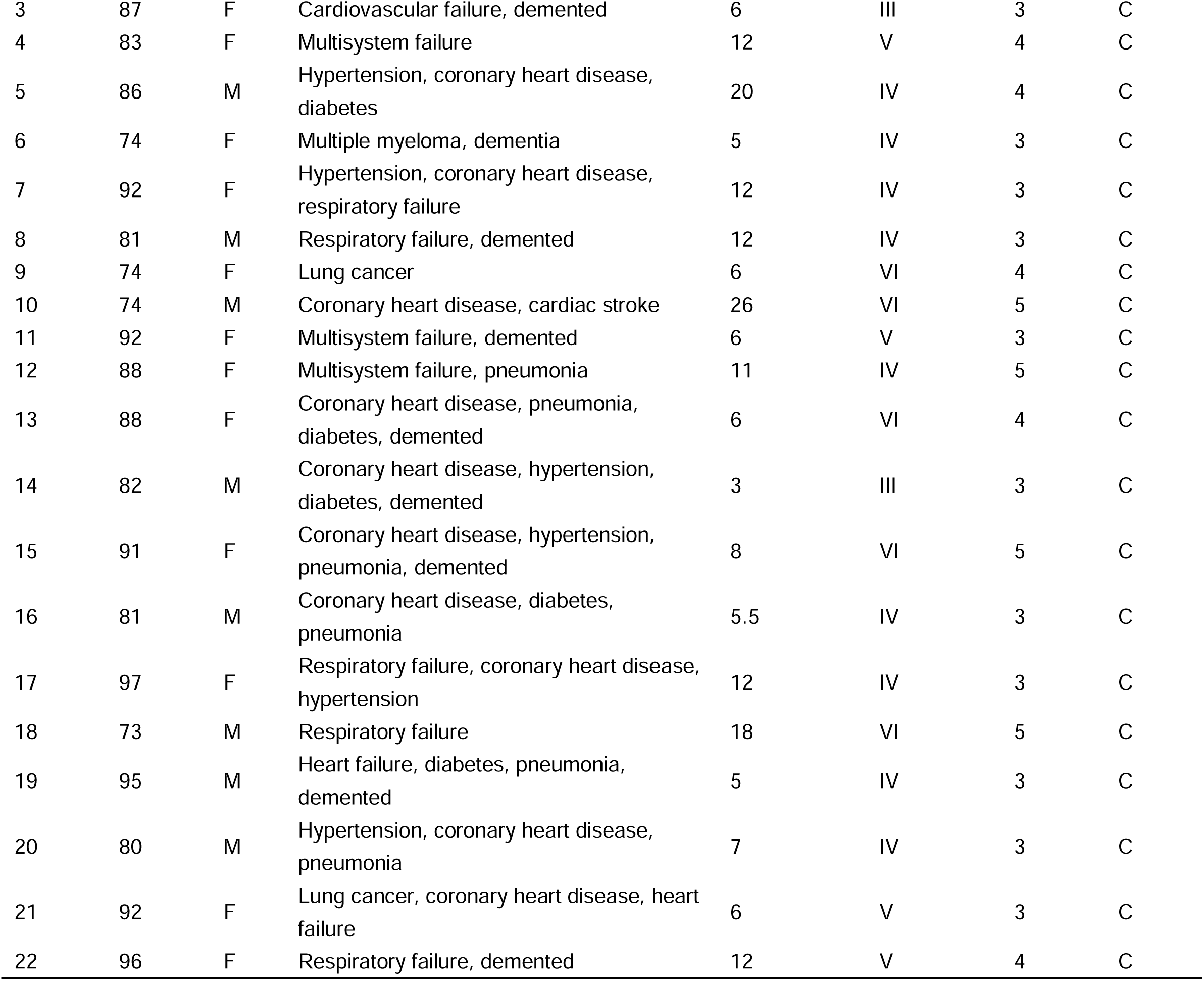
Information of the postmortem human brains used in the present study.

| Case number | Age (year) | Sex | Clinical diagnosis | Postmortem delay (hours) | Braak NFT stage | Thal Aβ phase | CERAD score |
| --- | --- | --- | --- | --- | --- | --- | --- |
| 1 | 75 | M | Multisystem failure | 5 | VI | 4 | C |
| 2 | 88 | F | Hypotension, heart disease, demented | 9 | V | 4 | C |
| 3 | 87 | F | Cardiovascular failure, demented | 6 | III | 3 | C |
| 4 | 83 | F | Multisystem failure | 12 | V | 4 | C |
| 5 | 86 | M | Hypertension, coronary heart disease, diabetes | 20 | IV | 4 | C |
| 6 | 74 | F | Multiple myeloma, dementia | 5 | IV | 3 | C |
| 7 | 92 | F | Hypertension, coronary heart disease, respiratory failure | 12 | IV | 3 | C |
| 8 | 81 | M | Respiratory failure, demented | 12 | IV | 3 | C |
| 9 | 74 | F | Lung cancer | 6 | VI | 4 | C |
| 10 | 74 | M | Coronary heart disease, cardiac stroke | 26 | VI | 5 | C |
| 11 | 92 | F | Multisystem failure, demented | 6 | V | 3 | C |
| 12 | 88 | F | Multisystem failure, pneumonia | 11 | IV | 5 | C |
| 13 | 88 | F | Coronary heart disease, pneumonia, diabetes, demented | 6 | VI | 4 | C |
| 14 | 82 | M | Coronary heart disease, hypertension, diabetes, demented | 3 | III | 3 | C |
| 15 | 91 | F | Coronary heart disease, hypertension, pneumonia, demented | 8 | VI | 5 | C |
| 16 | 81 | M | Coronary heart disease, diabetes, pneumonia | 5.5 | IV | 3 | C |
| 17 | 97 | F | Respiratory failure, coronary heart disease, hypertension | 12 | IV | 3 | C |
| 18 | 73 | M | Respiratory failure | 18 | VI | 5 | C |
| 19 | 95 | M | Heart failure, diabetes, pneumonia, demented | 5 | IV | 3 | C |
| 20 | 80 | M | Hypertension, coronary heart disease, pneumonia | 7 | IV | 3 | C |
| 21 | 92 | F | Lung cancer, coronary heart disease, heart failure | 6 | V | 3 | C |
| 22 | 96 | F | Respiratory failure, demented | 12 | V | 4 | C |

### Immunohistochemistry and immunofluorescence

Frozen and paraffin sections were immunohistochemically stained with the avidin-biotin complex (ABC) method with a set of primary antibodies diluted at optimal concentrations according to pilot experiments (see Table 2 for detailed information). The mouse anti-Aβ 6E10, mouse anti-pTau AT8, rabbit anti-sortilin CT and rabbit β-secretase 1 (BACE1) antibodies were consistently used for comparative assessment of immunolabeling between consecutive paraffin sections, with additional antibodies primarily used in fluorescent co-labeling experiments. The immunolabeling protocol was detailed in our recent studies (Hu et al., 2017; Jiang et al., 2022; Tu et al., 2020). Specifically, the paraffin sections were dewaxed in xylene and rehydrated through descending ethanol solutions, followed by antigen retrieval in ethylenediaminetetraacetic acid (EDTA, 1 mM, pH 8) at 98 °C for 10 minutes. For Aβ immunolabeling, the sections were additionally treated with 90% formic acid at room temperature for 10 minutes. Hematoxylin counterstain was applied on some immunolabeled sections for neuroanatomical orientation.

**Table 2:** Antibodies and key detection reagents used in the present study

| REAGENT or RESOURCE | SOURCE | IDENTIFIER |
| --- | --- | --- |
| Mouse monoclonal anti-Aβ 6E10 | Biologend | Cat# 803001; RRID:AB_2564653 |
| Mouse monoclonal anti-pTau AT8, | Invitrogen | Cat# MN1020; RRID:AB_223647 |
| Rabbit polyclonal anti-sortilin | Invitrogen | Cat# PA5-19481; RRID:AB_10986596 |
| Rabbit polyclonal anti-sortilin | Abcam | Cat# ab16640; RRID:AB_2192606 |
| Rabbit monoclonal anti-BACE1 | Abcam | Cat# ab183612; RRID:AB_3094757 |
| Goat polyclonal anti-Sortilin | R&D Systems | Cat# AF3154; RRID:AB_2286389 |
| Mouse monoclonal anti-MAP2 | Sigma-Aldrich | Cat# M9942; RRID:AB_477256 |
| Mouse monoclonal anti-GFAP | Cell Signaling Tech | Cat# 3670S; RRID:AB_561049 |
| Mouse monoclonal anti-Iba 1 | Proteintech | Cat# 66827-1-Ig; RRID:AB_2882170 |
| Alexa Fluor 488 donkey anti-rabbit IgG (H + L) | Invitrogen | Cat# A-21206; RRID: AB_2535792 |
| Alexa Fluor 594 donkey anti-rabbit IgG (H + L) | Invitrogen | Cat# A-21207; RRID:AB_141637 |
| Alexa Fluor 488 donkey anti-mouse IgG (H+L) | Invitrogen | Cat# A-21202; RRID:AB_141607 |
| Alexa Fluor 594 donkey anti-mouse IgG (H+L) | Invitrogen | Cat# A-21203; RRID:AB_141633 |
| Horse anti-Mouse/Rabbit/Goat IgG antibody (H+L) (Universal Pan-Specific), Biotinylated | Vector Laboratories | Cat# BA-1300; RRID:AB_2336188 |
| Tyramide signal amplification (TSA) | Thermo Fisher Scientific | Cat# T20922 and Cat# T20925 |
| Ready-to-Use Amylo-Glo kit | Biosensis Pty Ltd. | Cat#TR-300-AG, |
| Selective amyloid probe DANIR 8c | Cui MC | Fu et al., 2016 |
| Selective tangle probe TNIR7-1A | Cui MC | Xie et al., 2022 |

Double immunofluorescence was carried out using individual pairs of primary antibodies raised in different animal species (Table 2). The pretreatments of paraffin sections were the same denoted above, with the immunolabeling visualized using a proper combination of the Alexa Fluor® 488 and Alexa Fluor®594 conjugated donkey anti-mouse, anti-rabbit or anti-goat secondary antibodies (1:100, Invitrogen, Carlsbad, CA, United States). In a few settings, two primary antibodies raised in rabbits (i.e., two sortilin CT antibodies and rabbit anti-BACE1 vs. rabbit anti-sortilin antibodies) were used for double immunofluorescence, which was carried out with the tyramide signal amplification (TSA) method using commercial kits following manufacturer’s instruction (Cat# T20922 and Cat# T20925, ThermoFisher Scientific). This method involves the use of polymer-horseradish peroxidase (HRP) conjugated secondary antibodies that activate the fluorescently-tagged tyramide substrates in the tissue. The binding of the first antibody is then removed by heating (100 °C, 25 min, in phosphate-buffered saline), allowing a secondary round of primary antibody binding (regardless of its species origin) and fluorescent labeling. After the immunofluorescent labeling, the sections were briefly incubated in 0.1% Sudan black to block autofluorescence, then coverslipped with the VECTASHIELD Vibrance® antifade mounting medium containing the nuclear dye 4’,6-diamidino-2-phenylindole (DAPI) (Cat# H-1800, Vector Laboratories). In other experimental settings, double immunofluorescence was combined with counterstain of the Amylo-Glo fluorescent dye, which were also treated with Sudan black before mounting with a regular antifading mounting medium without DAPI (Cat#H-1900, Vector Laboratories). Immunofluorescent experiments were also carried out using one primary antibody, followed by counterstain with a fluorescent trace and DAPI (see below).

### Brain slice clearance, double immunofluorescent labeling and imaging

Double immunofluorescent labeling and 3D imaging of sorfra and Aβ plaques in the cerebral neocortex of an AD human brain (Braak stage V tau pathology and Thal phase 4 Aβ pathology) was carried out using the tissue clearance technology through a customized service (Nuohai Life Science Co., Ltd, Shanghai, China), following an established protocol (Chen et al., 2020). A tissue block (5×8.5×1 mm^3^) was dissected out at the superior temporal gyrus from the brain slice fixed in formalin for 1 week, with the slice further fixed for 10 hrs. The slice was cleared over 10 days in multiple replacements of solutions containing permabilization and dilapidation reagents. After the slice became transparent and treated with formic acid, it was incubated with refreshed (every two days) buffers containing mouse anti-Aβ antibody 6E10 (1:500) and rabbit anti-sortilin CT antibody (ab16640, 1:500) for 10 days, and subsequently with Alexa Fluor® 488 and Alexa Fluor®594 conjugated donkey anti-mouse and anti-rabbit secondary antibodies for 10 days. The whole slice was first scanned at a low resolution (3.3×3.3×7 μm^3^) on the Nuohai LS18 imaging system. A suitable region with a vascular landmark was identified, and scan-imaged at a high resolution (0.5×0.5×1.4 μm^3^). Vascular profiles were rendered in the 3D image (Chen et al., 2020).

### Histological staining with fluorescent amyloid and tangle tracers

Three fluorescent tracers were used in the present study to observe amyloid and tangle binding in brain sections. The Amylo-Glo dye stains β-amyloid deposits as well as mature and ghost tangles in bright-blue fluorescence (Yang et al., 2024), a ready-to-use kit of this reagent was obtained commercially and used according to manufacturer’s instruction (Cat#TR-300-AG, Biosensis Pty Ltd. SA5031, Australia). DANIR 8c is a recently developed red fluorescent probe for Aβ imaging (Fu et al., 2016), and TNIR7-1A is a high-performance near-infrared probe detecting neurofibrillary tangles *in vitro* and *in vivo* (Xie et al., 2022). These tracers were used in combination with double (Amylo-Glo and two primary antibodies) and single (DANIR 8c or TNIR7-1A with one primary antibody) antibody immunofluorescence. Briefly, following antibody immunolabeling, the paraffin sections were counterstained with Amylo-Glo, DANIR 8c or TNIR7-1A, and differentiated in 50% ethanol under microscopic monitoring (Fu et al., 2016; Xie et al., 2022; Yang et al., 2024). Strong acids can break down the β-pleated sheets and eliminates histological amyloid staining in tissue sections (Cammarata, 1990; Sun et al., 2002). Therefore, except for a specially designed experiment to test whether sorfra deposits are amyloid, all the sections used for Amylo-Glo, DANIR 8c or TNIR7-1A stain were not pretreated with formic acid.

### Gallyas and Bielschowsky silver stains

Gallyas silver stain was carried out according to an early physical development protocol (Gallyas, 1971). Briefly, paraffin sections were dewaxed and rehydrated, followed by incubations in 5% sodium periodate for 30 min and the “zilver jodide oplossing” solution containing sodium hydroxide, potassium iodide and silver nitrate for 30 min. The silver staining was precipitated in a developer [10:3:7 mixer of solution A (5% sodium carbonate), B (0.2% ammonium nitrate, 0.2% silver nitrate and 1% silicotungstic acid) and C (0.2% ammonium nitrate, 0.2% silver nitrate, 1% silicotungstic acid and 0.75% formalin) under microscopical monitoring, washed in de-ionized water (DW) and fixed in sodium thiosulphate (5%).

Gros’ Modified Bielschowsky’ silver stain was carried out on paraffin sections after dewaxation. The sections were brought into DW shortly and sensitized in freshly prepared 20% silver nitrate for 20 min. and placed in DW again. The ammoniacal silver solution was prepared by adding ammonia drop-wise into 20% silver nitrate, which was mixed until the precipitate dissolved and the solution became clear. Th sections were then impregnated in the dark in ammoniacal silver solution for approximately 15 min, until the neuronal fiber bundles and white matter stained dark and the background appeared tan. The sections were then processed in a developer (2.5 ml of citric acid, 2 ml of 40% formalin, one drop of nitric acid, and 95.5 ml of tap water), washed in DW, and fixed with sodium thiosulphate.

### Western blot

Western blot was used to appraise a new sortilin CT antibody (PA5-19481, Invitrogen) relative to the antibodies raised against the extracellular domain (AF3154, R&D Systems) and CT (ab16640, Abcam) of sortilin protein characterized in our earlier reports (Hu et al., 2017; Zhou et al., 2018). Temporal neocortical lysate was prepared from an AD brain and homogenized in 10 (w/v) radioimmuno-precipitation (RIPA) buffer containing a cocktail of proteinase inhibitors (Beyotime, China, #P0013B). The extract was centrifuged at 15,000 rpm for 15 min at 4 °C, with the supernatants collected and assayed for protein concentration (Bio-Rad Laboratories, Hercules, CA, USA). The sample was loaded into a furrow of 10% SDS-PAGE gel modeled with a teethless comb, separated by electrophoresis and transferred onto Trans-Blot pure nitrocellulose membranes. The membrane was cut into longitudinal stripes, which were immunoblotted separately with three sortilin antibodies, using horseradish peroxidase (HRP)-conjugated secondary antibodies and the PierceTM ECL-Plus Western Blotting Substrate detection kit to visualize the bound products (Thermo Fisher Scientific, MA, USA, #32132), with β-actin served as the internal loading control.

### Image acquisition

Bright-field immunolabeling and silver stain sections were scan-imaged on a Motic-Olympus microscope imaging system (Motic China Group Co. Ltd., Wuhan, Hubei, China). Images were examined and extracted with the Motic viewer software (Motic Digital Slide Assistant System Lite 2.0, Motic China Group Co. Ltd.). Fluorescently labeled sections were imaged with a Keyence KV-8000 imaging system (Keyence Corporation, Osaka, Japan) using a Z-stack setting of 1 µm scanning depth. The images were examined with the BZ-X800 Wide Image Viewer (Keyence Corporation, Osaka, Japan), with the whole image or selected regions exported, and further edited with the BZ-X800 Analyzer (Keyence Corporation, Osaka, Japan), including the preparation of overlapped and channel-separate micrographs, and insertion of scale bar or captured local areas.

### Quantitative image analysis and data processing

Areas in a size of 1000×1000 pixels were cropped from the original scanned images and saved as individual high-resolution files (tiff format) for quantification with the Image J software. Values of relative area (% of occupied relative to total area) were reported in the measurements of plaque profiles (sorfra, Aβ and amyloid tracer labeling). Optic densities were reported in the measurements of pTau and BACE1 immunolabeling, or Gallyas silver stained profiles. An identical optic density threshold was set for selecting the positive profiles among all the cropped micrographs of the same type of labeling. For sorfra plaque labeling, a low-limit size cutoff (100 μm) was used to eliminate the effect of sortilin labeled neuronal somata. The relative area and density readouts were normalized to the highest value in the same set of data, yielding comparable percentile levels (% of maximum) to allow correlative analysis. For correlative densitometry of sorfra and Aβ/amyloid labeling among individual plaque profiles, optic densities were reported with a threshold cutoff, measured in the areas outside the tissue section.

For each type of correlative analyses, stained sections from 3-4 brains were used. The microscopic zones planned for correlative analysis of sorfra plaque and pTau burdens were cropped randomly by referring to the X and Y coordinates of the double immunofluorescent image, with the zones fallen in the cortical gray matter considered effective (Ai et al., 2021; Li et al., 2023). The microscopic zones planned for correlative analysis of sorfra, Aβ and Gallyas silver labeling along the same sulcal to gyral transition were first set in the sorfra/Aβ double immunofluorescent image by visual judgment. Specifically, three zones were selected in the cortex around the sulcal bottom (“valley”), three zones around the middle portion of the sulcal wall, and three zones around corresponding gyral surface (“hilltop”). The fluorescent image with the zone map was transferred into the Photoshop file of silver-stained consecutive section as a separate layer, with the two images aligned to each other in reference to vascular landmarks. The frames of the measurement zones set before were copied as a template layer, with the fluorescent image *per se* deleted, allowing the selection of a corresponding set of sampling zones in the silver stain image for densitometry as described above. Microscopical zones planned for densitometry of sorfra relative to Aβ or amyloid tracer labeling were set in the hippocampal and cortical areas with a large proportion of colocalized plaque profiles. The cropped image fields were analyzed with Image J, reporting the optic densities of two types of deposits at individual plaques using a method reported earlier (Zhang et al., 2009).

### Statistical analysis and figure preparation

Correlative analyses were carried out between the normalized percentile values of two or three variables using the Graphpad 10 software (GraphPad Prism, San Diego, CA, United States). Dot graphs representing values derived from individual sampled zones or plaque profiles were plotted. Pearson correlation analyses were carried out to report the *P* value and coefficient index, with the minimal level of significant difference set at *P*<0.05. Figures were prepared with Photoshop 2024 by assembling representative low magnification whole section images and cropped high resolution microscopic views, together with graphs from data analyses appropriately.

## RESULTS

### Sorfra pathology was comparably detected by two sortilin CT antibodies

We have used a rabbit antibody raised against the sortilin C-terminal (Abcam, ab16640) in previous studies of sorfra pathology (Hu et al., 2017; Jiang et al., 2022; Tu et al., 2020; Zhou et al., 2018). Cross-validation of this lesion with additional antibodies is important for establishing its authenticity and robustness as a novel AD neuropathology. We noticed that a similar sortilin CT antibody (Invitrogen, PA5-19481) has become available, and were keen to replicate the sorfra pathology using this new antibody product (Figure 1A).

**Figure 1.**
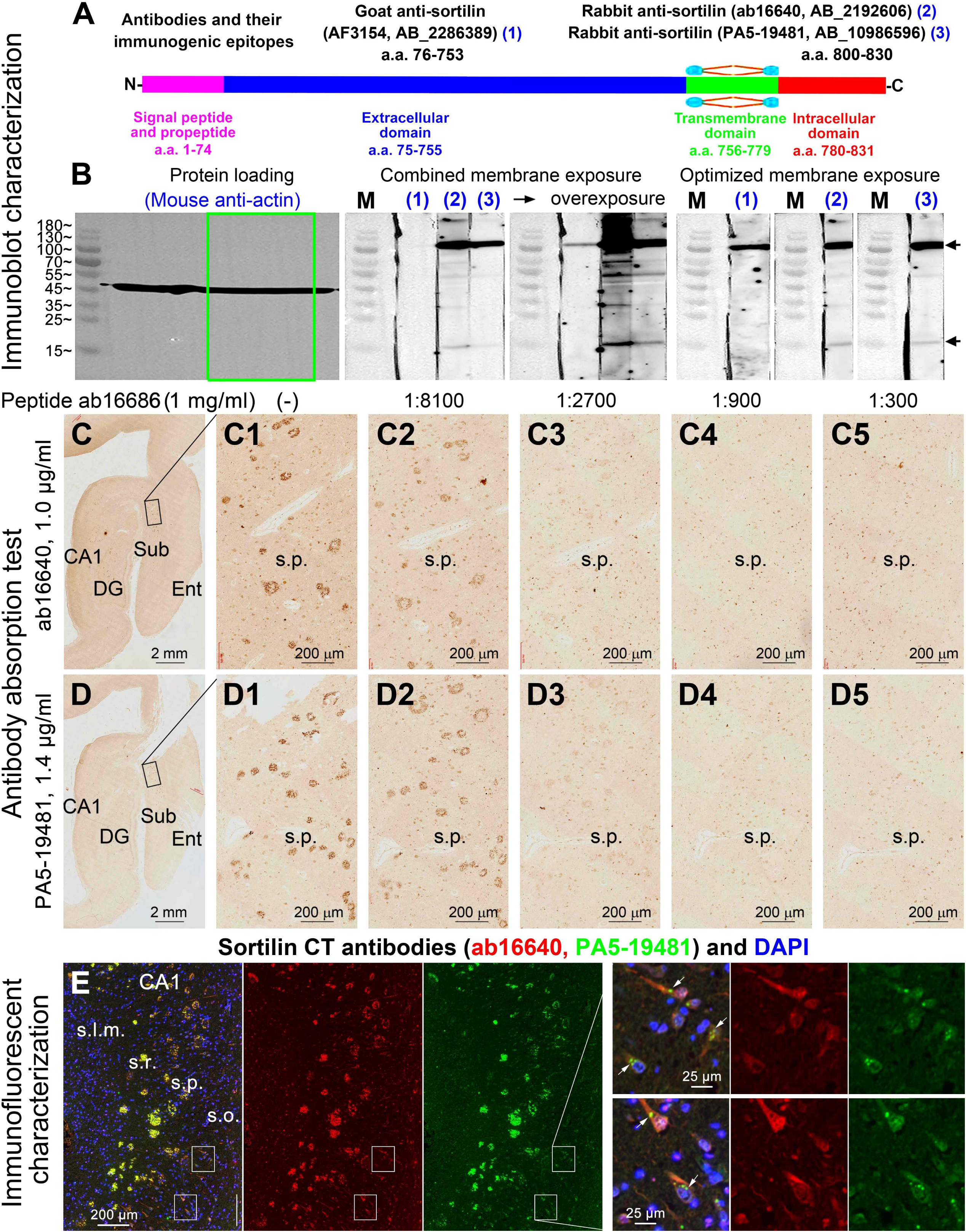
Western blot and immunohistochemical characterization of sortilin C-terminal antibodies for detecting sorfra pathology. **(A)** Structure of the sortilin protein and information about the antibodies and their immunogenic epitopes. **(B)** Immunoblot images showing the signal of the internal loading control β-actin and sortilin products detected by the three antibodies. Cortical lysate is filled into a furrow-shaped space in 10% SDS-PAGE gel, with the membrane and longitudinally cut strips (marked with a green outline) used for immunoblotting and reblotting. The full-length sortilin protein (100 kDa) is detected by the three antibodies, while the two C-terminal antibodies also detect bands migrated at ∼15 kDa and ∼55 kDa. **(C, C1-C5 and D, D1-D5)** The immunolabeling of the two C-terminal antibodies is diminished in the presence of the synthetic immunogenic peptide in a dose-dependent manner. **(E)** Overlapped immunofluorescent labeling of the two C-terminal antibodies in sorfra plaques, neuronal somata and dendrites, and intraneuronal inclusions (pointed by arrows) related to granulovacuolar degeneration. Experimental design, panel arrangement, antibody use and fluorescence indicators and scale bars for micrographic panels are as labeled. CA1: cornu Ammonis area 1, DG: dentate gyrus, Ent: entorhinal cortex, Sub: subiculum; s.o.: stratum oriens, s.p.: stratum pyramidale, s.r.: stratum radiatum, s.l.m.: stratum lacunosum-moleculare.

In western blot with an unequivocally identical loading of the cortical lysate from an AD human brain, the two sortilin CT antibodies immunoblotted protein products migrated at similar molecular weight locations, corresponding to the ∼100 kDa full-length sortilin and the ∼15 kDa fragment, with another ∼55 kDa band also visible (Figure 1B). In comparison, the goat antibody raised against the sortilin extracellular domain (R&D Systems, AF3154) essentially detected the full-length protein. The immunolabeling of both CT antibodies could be absorbed by the synthetic antigenic peptide (a.a. 800-830) in a dose-dependent manner (Figure 1C-C5, 1D-D5), and appeared to be completely colocalized in double immunofluorescence (Figure 1E). At closer examination, both CT antibodies visualized morphologically heterogenous extracellular deposition in paraffin and cryostat sections, ranging from scanty fibril clusters approximately in a size of a neuron, to heavy massed plaques mostly in a diameter from 50-100 μm (Figure 1C1, D1, E; Supplemental Figure 1). Some of the small fibril deposits appeared to align with the apical dendrites of cortical and hippocampal pyramidal neurons (Supplemental Figure 1A1). As with our earlier observations (Hu et al., 2017; Shi et al., 2020; Jiang et al., 2022), both antibodies could visualize neuronal somata and dendrites in the brains or brain areas without or with a lesser extent of sorfra plaque lesions. In addition, these antibodies displayed intraneuronal inclusions related to granulovacuolar degeneration (GVD) (Figure 1C1, D1, E; Supplemental Figure 1A). The PA5-19481 antibody was primarily used in the current study considering a cost-effective benefit (worked at higher dilutions).

### Sorfra, pTau and tangle pathologies developed in parallel over cerebral regions

Consecutive paraffin sections were prepared from recently banked brains with initially scored Aβ pathology from Thal phases 3-5 and tau pathology from Braak stages III-VI. The distribution and burden of Aβ, sorfra and pTau immunolabeling, and Gallyas stained tangle profiles, were assessed microscopically and quantitatively. Sorfra plaques were distributed over the cerebral regions in a progressive manner, which was generally in parallel with the progression of pTau and silver tangle labeling, but not Aβ deposition. Thus, in the brains with tauopathy at Braak stages III-IV, a parallel progression of sorfra, pTau and tangle profiles was seen from the limbic structures to the isocortex (Figure 2; Supplemental Figures 2, 3). In the occipital cortex, a parallelly spreading distribution of sorfra, pTau and tangle labeling from area 18 into area 17 was observed while comparing sections from the brains with tauopathy from Braak stage IV to VI (Supplemental Figures 3, 4, 5, 6). However, Aβ plaques and BACE1 labeled dystrophic neurites were present in the primary and secondary visual cortices in these brains in a largely comparable manner (Supplemental Figures 4, 5, 6). Notably, sorfra, pTau and tangle profiles could appear more denser in one gyrus relative to another in the same cortical region, sometimes even between two sides of the cortex bordering a single sulcus or fissure (Figure 2; Supplemental Figures 7, 8, 9).

**Figure 2.**
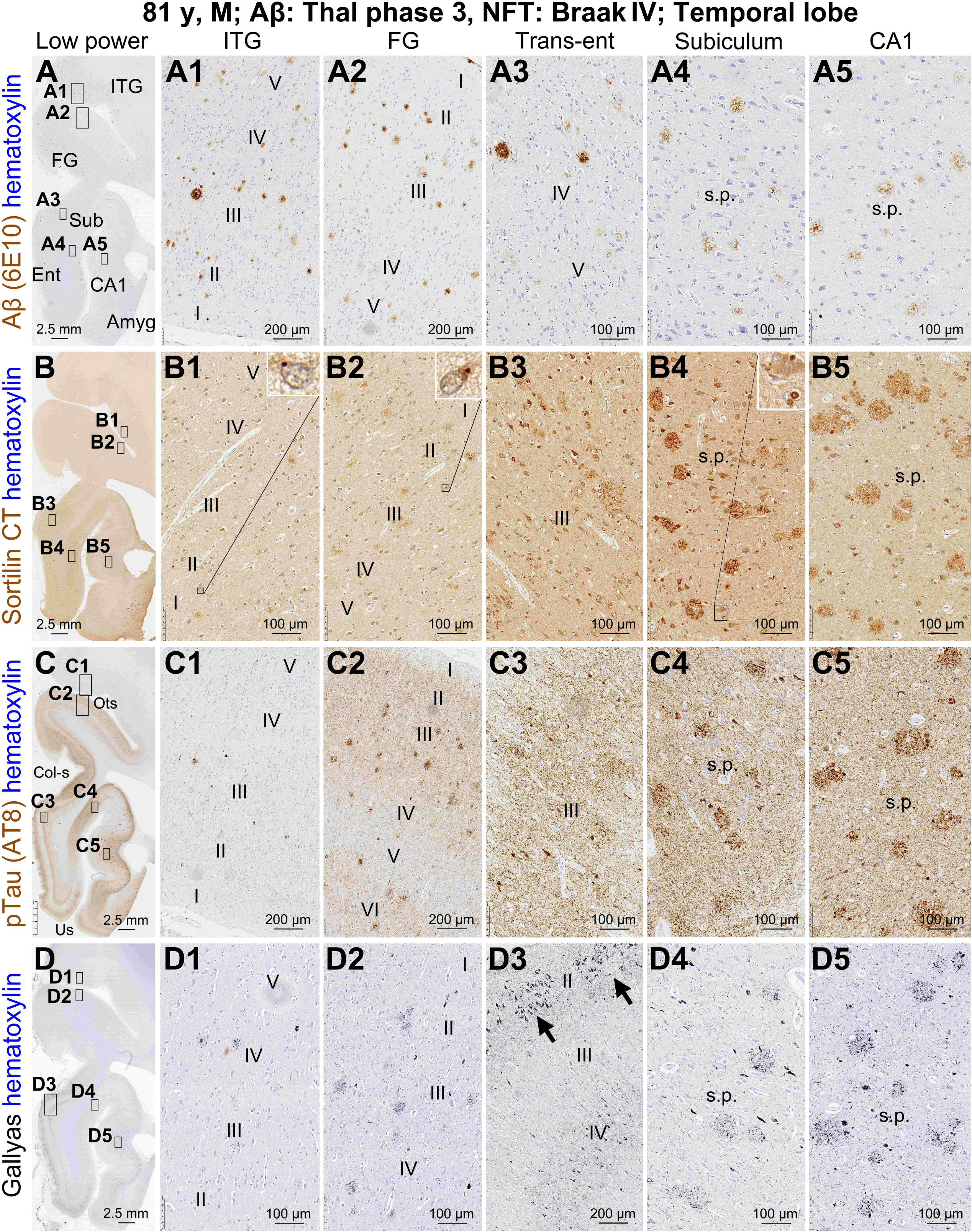
Parallel regional distribution of sorfra, pTau and Gallyas tangle pathologies, but not Aβ deposition, in the temporal lobe in a probable AD case. Low magnification maps of the labelings are shown on the left **(A, B, C, D)**, with boxed areas enlarged as indicated. The overall amount and labeling intensity of the Aβ plaques appear to be greater in the temporal neocortex relative to entorhinal cortex (Ent) and hippocampal formation **(A1, A2, A3, A4, A5)**, and appears comparable between the inferior temporal gyrus (ITG) and fusiform gyrus (FG) across the occipitotemporal sulcus (Ots). Sorfra plaques **(B1, B2, B3, B4, B5)**, pTau labeling **(C1, C2, C3, C4, C5)** and silver stained tangles **(D1, D2, D3, D4, D5)** exhibit a differential distribution over the regions opposite to the trend of Aβ. Plaque-like structures in sorfra, pTau and silver labeling are more distinctly present in subiculum (Sub)/CA1 relative to the entorhinal, transentorhinal (Trans-ent) and temporal neocortical areas. The overall amount of plaque-like profiles and their labeling intensity are greater in the fusiform gyrus than in the inferior temporal gyrus **(B1, B2, C1, C2, D1, D2)**. Note that Gallyas stain reveals a large amount of ghost tangles in the entorhinal and transentorhinal cortex, especially in the layer II cell islands **(D3,** arrows), whereas pTau labeling reveals a much smaller number of neurons in these areas **(C3)**. Also note the intraneuronal inclusions labeled by the sortilin C-terminal antibody (**B1, B2, B4**, inserts). Additional abbreviation: Amyg: amygdala, Col-s: collateral sulcus, Us: uncal sulcus. I-VI: cortical layers, s.p.: stratum pyramidale. Scale bars are as indicated.

An impressive observation was the differential occurrence of sorfra plaques together with pTau and silver tangle staining in the cortex as moving from the sulcal bottom or valley to the gyral surface. Specifically, there existed a high to low gradient in the amount of sorfra plaques, pTau labeled neuronal and neuritic plaques, and silver-stained neuritic plaques, tangles and neuropil threads in the cortical grey matter around the sulcal valley relative to the middle segment of the sulcal banks, and further to the corresponding gyral hilltop. However, the overall amount or density of Aβ plaques appeared otherwise comparable or less differentiated from the sulcal to gyral cortex (Supplemental Figures 2, 3, 7, 8, 9).

Temporal and frontal lobe sections selected from three brains were processed in the same batch for sorfra and pTau double immunofluorescence for a correlative analysis of the lesions in randomly sampled cortical zones (Figure 3A, A1, A2). Additional consecutive sections were immunofluorescently stained for Aβ and sorfra, and histologically stained for tangles with the Gallyas method, for a correlative analysis of the lesions in reference to the transition from sulcal valley to gyral hilltop (Figure 3B, B1, B2; C, C1, C2). A positive correlation (P<0.0001, r=0.52) was found between the normalized fractional areas (% of maximum) of sorfra plaques and normalized specific densities (% of maximum) of pTau labeled profiles among 181 sampled cortical zones (1000×1000 pixels in size) (Figure 3A, A1, A2, D). Among 21 sets of cortical zones (9 zones *per* set) quantitatively analyzed, there was a progressive decrease in the fractional areas of sorfra plaques and the specific densities of silver-stained tangle profiles from the sulcal valley to gyral hilltop areas. Person analysis indicated a positive correlation between the sorfra and tangle variables (P=0.0001, r=0.95), with the trendline of the former displayed a sharper slope relative to the latter (Figure 3E). The values of Aβ plaques in the sampled zones showed a flat trendline as moving from the sulcal valley to gyral hilltop, with no correlation to either the sorfra (P=0.384, r=-0.33) or tangle (P=0.601, r=-0.20) variables (Figure 3E).

**Figure 3.**
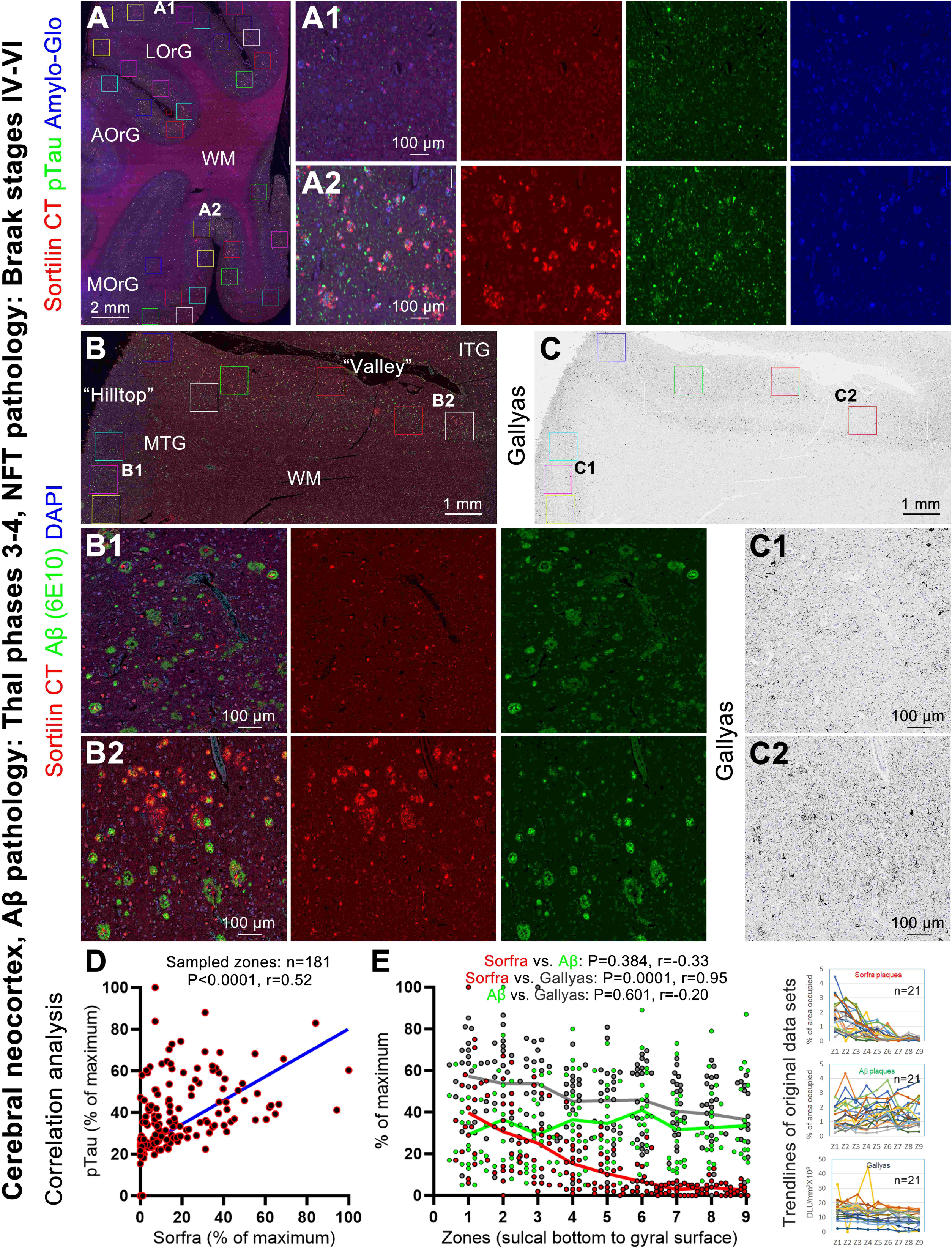
Correlative densitometry of sorfra, Aβ and neurofibrillary tangle pathologies in neocortical regions. **(A)** A representative image of sorfra, pTau and Amylo-Glo triple labeling in a frontal cortical section. Two zones located in the gyral hilltop (**A1**) and sulcal valley (**A2**), respectively, are enlarged to illustrate the regional difference in the overall amount of the labeled profile. (**B, C**) Representative images of sorfra and Aβ double immunofluorescence and Gallyas silver stain in consecutive paraffin sections over a temporal neocortical area, with enlarged panels (**B1, B2, C1, C2**) showing the sorfra/Aβ colocalization pattern and the overall amount of labeled profile in the cortex along the sulcal valley to gyral hilltop transition. In the first experimental setting, the fractional areas (% of occupied relative to total area) of sorfra plaques and specific densities (with subtraction of background density measured outside the section area) of pTau labeled profiles among randomly sampled zones are correlatively analyzed following normalization to the highest readout in each data set (expressed as % of maximum), indicating a positive correlation between the two measurements (**D**). In the second experimental setting, correlative densitometry for sorfra plaques, Aβ plaques and Gallyas labeled profiles are carried out in 9 cortical zones moving from the sulcal valley to gyral hilltop along each transition. Quantitative data from 21 sets of this regional transition are shown in panel (**E**). There is a parallel and corelated decline of the sorfra plaques and Gallyas silver-labeled profiles from the valley to hilltop areas, whereas this trend is not reflected in the Aβ measurement. AOrG: anterior orbit gyrus, MOrG: medial orbit gyrus, LorG: lateral orbit gyrus, MTG: medial temporal gyrus, ITG: inferior temporal gyrus, WM: white matter. Scale bars are as indicated.

### Sorfra plaques and neuritic plaques were anatomically matchable

Neuronal tangle formation is a continuous process conventionally divided into pretangle, mature tangle and ghost tangle stages based on morphological assessment with pTau immunolabeling and various histological stains (Moloney et al., 2021). We first determined the morphological matchiness between Aβ, sorfra and neuritic plaques using consecutive sections alternatively stained with immunohistochemistry, and Gallyas and Bielschowsky silver stains (Figure 4A, B, C, D, E, Supplemental Figure 10). In temporal lobe sections from a brain with Thal phase III Aβ pathology and Braak stage IV NFT pathology, neuritic plaques in CA1 and subiculum were packed with argyrophilic dystrophic neurites against a lightly stained background in silver stains. Prominent deposition of sorfra fibrils occurred in these plaques, with isolated fibrils visible in some silver stained somata or large dendritic trunks (Figure 4A, A1, A2; C, C1, C2, D, D1, D2; E, E1, E). In comparison, the 6E10 antibody displayed light extracellular Aβ deposition in the plaques, with some spheric profiles also labeled, likely representing swollen presynaptic terminals (Figure 4B, B1, B2) (Zhang et al., 2009). The 6E10 antibody visualized compact Aβ plaques in the subiculum, entorhinal cortex and temporal neocortex, along with meningeal, subpial and vascular Aβ deposition. Some compact Aβ plaques had a dense core or appeared in a circular form. Despite a heavy Aβ deposition, there were no or only a few silver-stained neurites in these plaques. A slightly increased background-like silver and sorfra labeling was seen in these Aβ plaques, relative to the peri-plaque areas (Figure 4A1, B1, C1, D1 and E1, inserts). In another set of temporal lobe sections from a brain with Aβ pathology in phase 5 and NFT pathology at Braak stage IV, sorfra plaques and neuritic plaques (Bielschowsky stain) in the hippocampal formation were well matched in shape, size and contexture between consecutive sections, whereas only light Aβ labeling was seen in the plaques. Again, an opposite picture was seen in the temporal lobe cortex, wherein heavily labeled compact Aβ plaques contained no or only isolated sorfra deposits and silver stained neuritic processes (Supplemental Figure 11).

**Figure 4.**
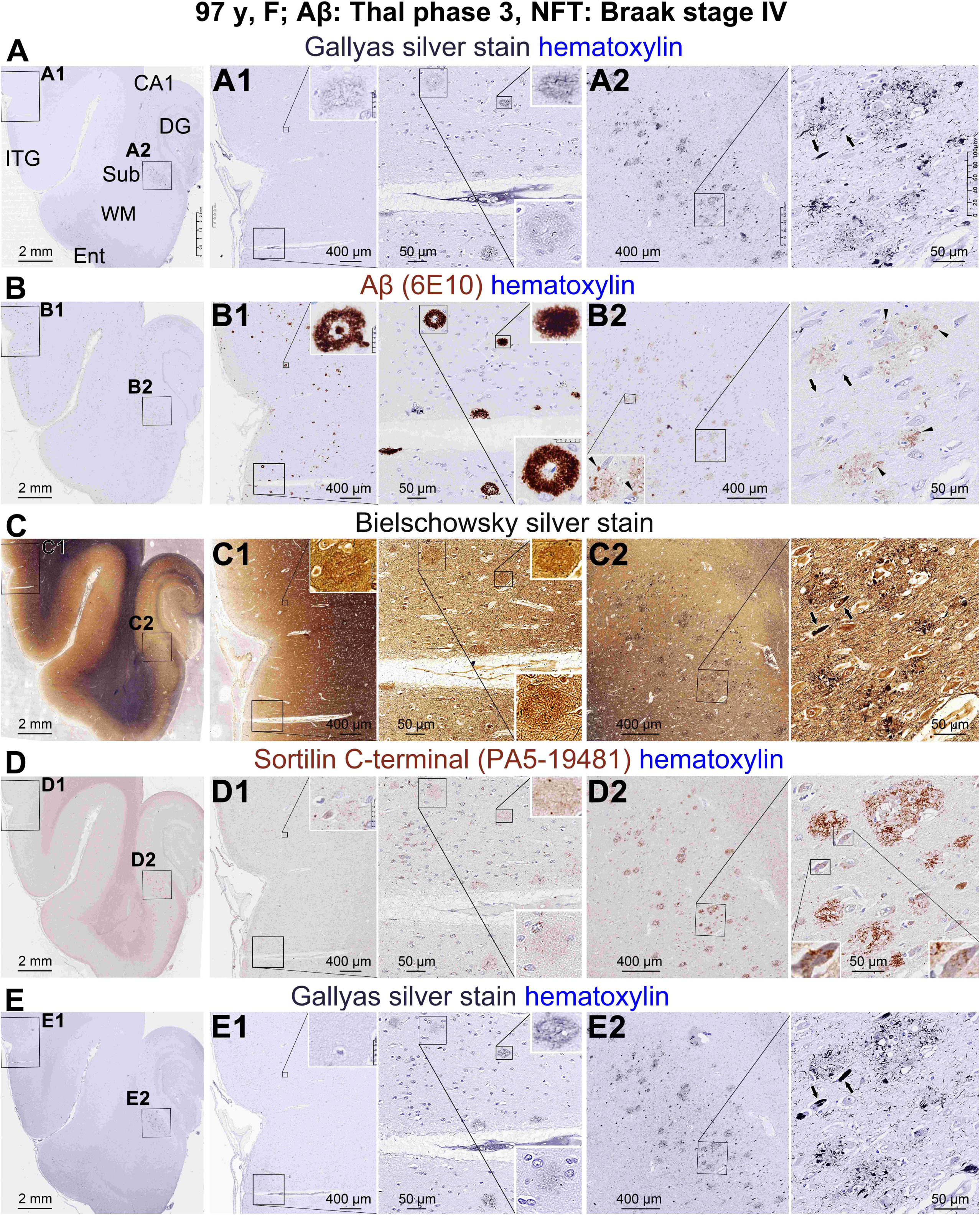
Assessment of the anatomical matchiness between Aβ, sorfra and neuritic plaques across consecutive temporal lobe paraffin sections. Information of the brain sample, staining methods and anatomical locations is provided, with low magnification maps (**A, B, C, D, E**) and enlarged views illustrated (**A1, A2, B1, B2, C1, C2, D1, D2, E1, E2**). Aβ plaques are more densely distributed in the temporal neocortical relative to entorhinal (Ent) areas and to the subiculum (Sub), CA sectors and dentate gyrus (DG). In a close view from the inferior temporal gyrus (ITG) (**A1, B1, C1, D1, E1**), heavily stained Aβ plaques are cored, massed or in circular shape (inserts); they contain few silver-stained neurites and exhibit slightly increased extracellular silver and sorfra labeling relative to plaque periphery. The plaque structures in the hippocampal CA1 and subicular areas are matchable in location, size and shape between Gallyas (**A2, E2**), Bielschowsky (**C2**) and sorfra (**D2**) staining preparations, whereas these plaques contain only light extracellular Aβ labeling (**B2**) along with some labeled swollen neurites (**B2**, pointed by arrowheads). The sorfra fibrils are packed in the extracellular space in these plaques, which are also visible in the tangle-bearing somal and dendritic profiles in the silver stains (right most panels, pointed by arrows). Scale bars are as indicated.

We further determined the morphological matchiness using same-section double and triple fluorescent labeling. The first approach was double immunofluorescence for pTau and sorfra combined with Amylo-Glo counterstain (Figure 5; Supplemental Figures 12, 13, 14); the latter visualizes both amyloid and tangle (Yang et al., 2024). In this triple fluorescent preparation of temporal lobe sections with late-stage AD pathologies, a co-evolving pattern of the labeled profiles was seen over the hippocampal and temporal cortical areas. In the hippocampal formation, the subiculum and prosubiculum showed a large amount of sorfra plaques, which tended to reduce as moving into the CA1 to CA3 sectors or towards the presubiculum and parasubiculum. The extent of Amylo-Glo staining was in parallel with the above cross-region transitional trend. pTau labeling appeared relatively light in the central area with dense sorfra plaques but became increased as moving to the border wherein the sorfra plaques started to reduce (Figure 5A; Supplemental Figures 12A). At high magnifications (Figure 5A1; Supplemental Figures 12A1, A2, A3; 13A1, A2), Amylo-Glo visualized neuronal somal profiles appearing as mature and ghost tangles with and without pTau co-labeling, respectively. Amylo-Glo stained dystrophic neurites with and without pTau co-labeling coexisted in the sorfra plaques. Moving peripherally to the area wherein sorfra plaques started to reduce, we saw pTau positive neurites with no or less Amylo-Glo staining coexisting with sorfra deposition. A similar co-evolving pattern could be seen in the transentorhinal area, wherein Amylo-Glo and sorfra labeling appeared heavy whereas pTau labeling appeared relatively light. As moving into the entorhinal cortex and the temporal neocortex, pTau labeling became more microscopically impressive (Figure 5A, Supplemental Figure 12A). At high magnification, Amylo-Glo stained amyloid plaques colocalized differentially with sorfra plaques (will be detailed further), while this dye also stained neuronal and neuritic profiles with and without pTau co-labeling as examined closely (Figure 5B; Supplemental Figure 12A4). In the temporal neocortex (Supplemental Figure 13A, A1, A2, A3) as well as other neocortical areas (Supplemental Figure 13B, C), neuritic processes labeled by pTau, Amylo-Glo or both co-existed with the sorfra plaques. However, the overall Amylo-Glo intensity was variable among the sorfra plaques, displaying a differential pattern regarding the distribution and intensity of Amylo-Glo relative to sorfra among individual plaque profiles, as addressed below.

**Figure 5.**
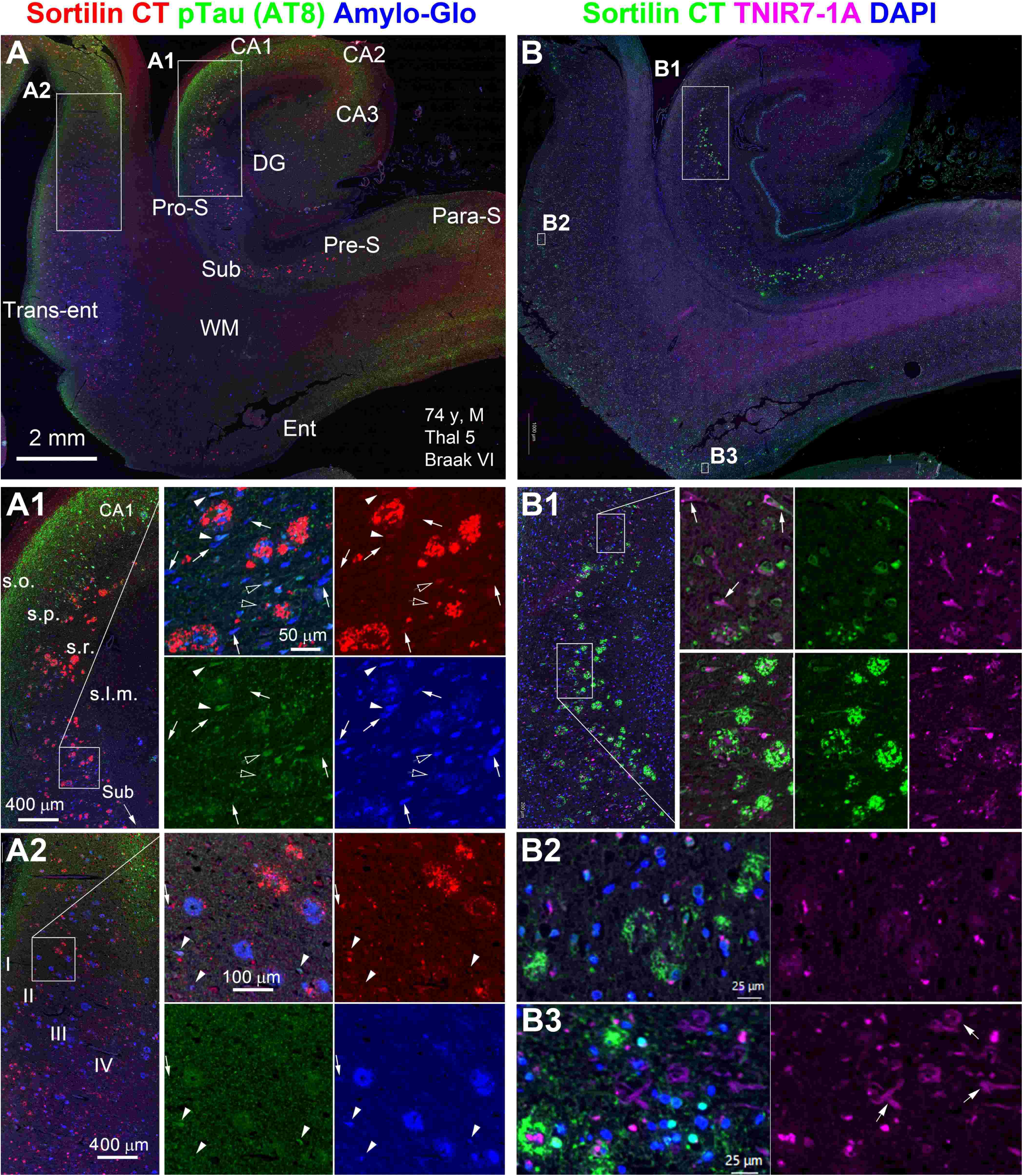
Double and triple fluorescent characterization of sorfra plaque colocalization with neurofibrillary tangles. Shown are adjacent temporal lobe paraffin sections from an AD human brain immunofluorescently labeled for sorfra and pTau with Amylo-Glo counterstain (**A, A1, A2**), and immunofluorescently labeled for sorfra and stained with a selective high-performance near-infrared tangle probe TNIR7-1A and DAPI (**B, B1, B2**), with the fluorescent indicators marked. A local transitional pattern is visible at low magnification in the sorfra/pTau/Amylo-Glo preparation in the subicular/CA area and the transentorhinal area (Trans-ent). Sorfra and Amylo-Glo labeling appear heavy in a middle zone and become lighter as moving into its periphery, whereas pTau immunofluorescence shows an opposite trend (**A**). In the hippocampal formation, Amylo-Glo stain displays mostly somal and neuritic profiles that are differentially colocalized with pTau immunofluorescence, including singly labeled ghost tangles (arrows) and double-labeled mature tangles (arrowheads). Clusters of the single and double labeled neurites are present in the sorfra plaques (**A1**). Mature-looking tangles (pTau/Amylo-Glo positive) show light sortilin labeling with inclusions (hollowed arrows). In the entorhinal cortex (Ent), pTau labeled neurites and Amylo-Glo stain are also present in the sorfra plaques (**A2**). A mixed sorfra and Amylo-Glo colocalization pattern is seen among the plaque profiles. In this case, it is difficult to judge whether the Amylo-Glo labeling reflects Aβ deposition, tangle-bearing neurites or both. In the sorfra/ TNIR7-1A preparation (B), the tangle probe displays neuritic clusters in both the hippocampal and cortical sorfra plaques (B1, B2). This selective probe displays somal profiles likely representing mature tangle and pretangle that exhibit light sortilin colocalization including intracellular granules (B1, pointed by arrows). This tracer also visualizes ghosts tangles especially abundant in the entorhinal cortex (B3, pointed by arrows). Additional abbreviations: CA1-3: subregions of cornu Ammonis, DG: dentate gyrus, Pro-S: prosubiculum, Pre-S: presubiculum, Para-S: parasubiculum, I-IV: cortical layers; WM: white matter, s.o.: stratum oriens, s.p.: stratum pyramidale; s.r.: stratum radiatum, s.l.m.: stratum lacunosum-moleculare. Scale bars are indicated.

Because Amylo-Glo stains both amyloid and tangle, it was sometimes difficult to distinguish whether the staining in a plaque was related to amyloid or tangle-bearing neuritic processes, especially in the cortical plaques (Figure 5A2; Supplemental Figures 12A4; 13). Therefore, we used a selective near-infrared tangle probe, TNIR7-1A, to examine the matchiness between sorfra and neuritic plaques (Xie et al., 2022). With this approach, sorfra deposits and tangle-like neuritic profiles were found to coexist at the plaque loci in the hippocampal formation and cortical regions. This tangle probe also labeled somal profiles morphologically characteristic of mature and ghost tangles. In addition, this tracer labeled GVD-like inclusions in neuronal somata likely at mature and pretangle stages, with the putative pretangle neurons also exhibited light sortilin labeling (Figure 5B, B1, B2, B3; Supplemental Figure 14B, B1, B2, B3).

The TNIR7-1A tracer appeared fairly potent to probe early tangle-like changes in neuronal somata and dendritic processes, therefore we further explored the early occurrence of sorfra deposits relative to TNIR7-1A labeling. In an aged human brain with relatively low burdens of Aβ and tau pathologies in the neocortex and hippocampus (Supplemental Figure 14A, B, C). TNIR7-1A labeling was seen in many cortical and hippocampal pyramidal neurons co-labeled by the sortilin CT antibody (Supplemental Figure 15D, D1, D2, D3, D4, D5). Sortilin labeled granular inclusions occurred inside the soma and dendrites, while some appeared to be outside the dendritic processes and deposited as small plaque-like structures (Supplemental Figure 14B, B2, B3). TNIR7-1A labeled dystrophic neurites were visible at these sparsely occurred, likely initially formed, extracellular sorfra plaques (Supplemental Figure 15D1, D2, D4).

### Sorfra and Aβ deposited as colocalized as well as separated plaques

As mentioned above, there existed a differential colocalization pattern between Amylo-Glo and sorfra labeling among the plaque profiles, with singly labeled profiles being the two extremes. This pointed to a heterogenous or co-evolving pattern of sorfra and Aβ deposition forming plaque structures. To address whether the application of Aβ antibodies could impact the effectiveness of microscopic detection of the plaques, we carried out double immunolabeling for Aβ and sorfra by using a single pair of antibodies and by simultaneously using the two rabbit sortilin CT antibodies combined with three mouse anti-Aβ antibodies. A differential Aβ and sorfra colocalization pattern, i.e., existence of colocalized as well as separated plaques, was observed in the above experimental settings without distinguishable difference (Supplemental Figure 16). Formic acid can break down the β-pleated sheet structure, thereby eliminating the binding of amyloidogenic proteins to amyloid dyes (Cammarata et al., 1990; Styren et al., 2000; Sun et al., 2002). On the other hand, this treatment enhances Aβ antibody labeling likely by increasing the exposure of antigen epitopes (Kitamoto et al., 1987). We carried out Aβ and sorfra double immunofluorescence with Amylo-Glo counterstain with and without formic acid pretreatment of the paraffin sections (Supplemental Figure 17). In the formic treated sections, a differential sorfra and Aβ colocalization pattern was seen among the plaque profiles, while the Amylo-Glo stain appeared largely background-like (Supplemental Figure 17A, C). In the formic untreated sections, a differential sorfra and Amylo-Glo colocalization pattern was observed, while the Aβ plaques were faintly labeled and visible by close examination (Supplemental Figure 17B, D).

Tissue clearance 3-D imaging technology was used to further verify that sorfra and Aβ can deposit as colocalized and separate plaques (Supplemental Figure 18). In a neocortical slice from an AD brain immunofluorescently labeled with sortilin CT (ab16640, Alexa Fluor®594 fluorescence) and Aβ (6E10, Fluor® 488 fluorescence) antibodies with vascular rendering, plaques appearing in green, yellow, orange and red were visible throughout the cortex in the image scanned at low resolution. In a local region scanned at high resolution, individual plaques exhibited the above spectrum of fluorescence in the 3-D image and across single scanning planes. Pure Aβ (green) plaques occurred densely near the cortical surface and could be related in part to meningeal and subpial deposition. Mixed (yellow and orange) and pure (red) sorfra plaques were largely located in the deeper cortical layers (Supplemental Video).

### Sorfra and Aβ densities were inversely correlated among the colocalized plaques

To understand the dynamics between sorfra and Aβ deposition among the colocalized plaques, densitometry was carried out using formic treated and untreated sections with sorfra/Aβ double immunofluorescence and Amylo-Glo counterstain (Figure 6A, B, C, D). Under the formic treated condition, the normalized values (% of maximum) of sorfra density were negatively correlated with that of Aβ density among the colocalized plaques measured over CA1 and subiculum (P<0.0001, r=-0.242, number of plaques n=306) and in the temporal neocortex (P<0.0001, r=-0.279, n=262) from two AD brains (Figure 6A1, A2). Under the formic untreated condition, a negative correlation between sorfra and Amylo-Glo densities was found among the plaques in CA1/subiculum (P=0.0005, r=-209, n=277) and in the temporal neocortex (P<0.0001, r=-503, n=233) (Figure 6C1, D1). Considering the amyloid/tangle dual staining issue of Amylo-Glo as noted above, sections were immunolabeled for sorfra and counterstained with a specific fluorescent Aβ probe, DANIR 8c (Fu et al., 2016), and DAPI (Figure 6E, F; Supplemental Figure 19). A negative correlation existed between sorfra and DANIR 8c densities among the colocalized plaques in the hippocampal formation (P<0.0001, r=-355, n=377) and temporal neocortex (P=0.005, r=-136, n=423) (Figure 6E1, F1). Sorfra positive and DANIR 8c negative as well as DANIR 8c positive and sorfra negative plaques were concurrently present in the sections (Figure 6E, F; Supplemental Figure 19).

**Figure 6.**
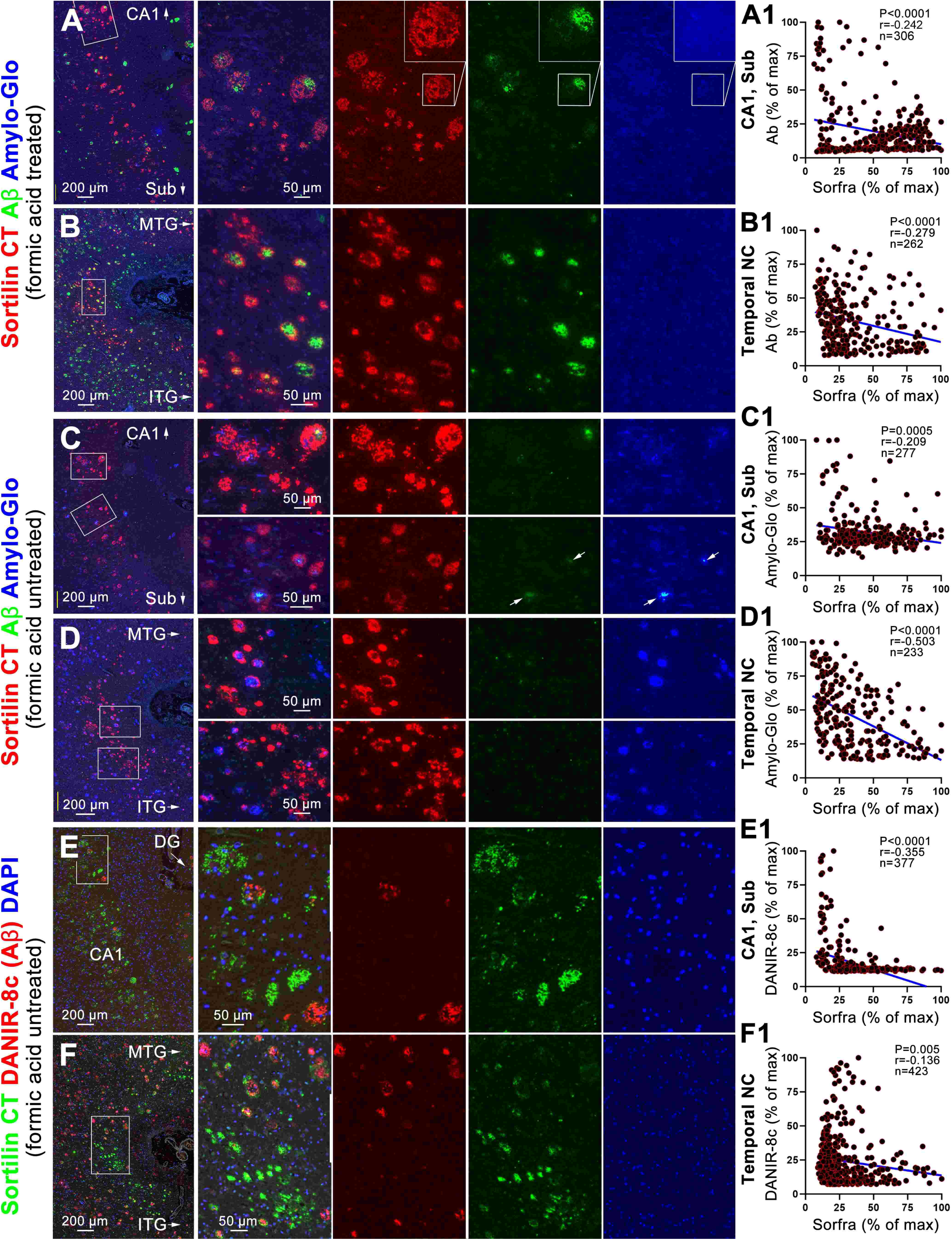
Correlative densitometric analyses of sorfra, Aβ and amyloid tracer labeling in hippocampal and neocortical plaques. Images are from temporal lobe paraffin sections immunofluorescently labeled for sorfra and Aβ followed by Amylo-Glo counterstain in conditions with (**A, B** and enlarged views) and without (**C, D** and enlarged views) formic acid pretreatment, and from sections immunofluorescently labeled for sorfra followed by counterstain with a specific amyloid probe, DANIR-8c (**E, F**). Densitometry is carried out using Image J in cropped microscopical zones (1000×1000 pixels) containing mostly the colocalized plaques with a few non-colocalized ones, with latter used to obtain the highest readouts (i.e., the maximum) of sorfra or amyloid labeling for normalization purpose. In all the above settings, a differential labeling of sorfra relative Aβ or the amyloid tracers exist among the plaque profiles (enlarged panels). The Amylo-Glo stain appears largely background-like in the condition of formic pretreatment, with some tangle-like profiles still visible at high magnification (A, insert). Without formic acid treatment, the Aβ immunolabeling is largely not visualized except for the sites with the heaviest amyloid load (enlarged panels in C, bleached in the image, pointed by arrows). In this preparation, the Amylo-Glo labeled profiles appear largely in somal and neuritic shapes in the subiculum (Sub) and CA1 areas, representing tangle-bearing structures (C1 and enlarged panels), while those in the temporal neocortex include plaques (with relatively larger size and stronger labeling) as well as tangle-bearing somata and neurites (visible by close examination). The right most panels show the densitometric dot graphs including results of statistical analysis. An inverse correlation exists for sorfra relative to Aβ densities (A1, B1), sorfra relative to Amylo-Glo densities (C1, D1) and sorfra relative to DANIR-8c densities (E1, F1) among the plaques quantified in the subiculum to CA1 region and in the neocortical regions including the inferior temporal gyrus (ITG), middle temporal gyrus (MTG) and others (shown in supplemental figures).

BACE1 is the obligatory enzyme initiating Aβ production through the amyloidogenic cleavage pathway of the β-amyloid precursor protein (APP) processing. This enzyme is enriched at axonal terminals under normal condition and elevated in presynaptic dystrophic neurites in Aβ plaques (Cai et al., 2010; Sadleir et al., 2016; R. Yan, 2017; Zhang et al., 2009). We further explored the relevance of sorfra deposition to the alteration of presynaptic dystrophic neurites expressing BACE1. In the visual cortical areas from brains with tauopathy advancing from Braak stages IV to VI, BACE1 labeled swollen axonal terminals occurred in both areas 17 and 18 in parallel with the presence of Aβ plaques (Supplemental Figures 5, 6, 7). In sorfra/pTau double immunofluorescence with Amylo-Glo counterstain, BACE1 labeled neuritic clusters frequently occurred in Amylo-Glo stained plaques in areas 17 and 18 in the brains with tauopathy not yet reached these areas (Supplemental Figure 20A, B, C). In the brains with Braak stage VI tauopathy, BACE1 labeled dystrophic neurites colocalized with sorfra and Amylo-Glo stained plaques in a differential manner. Thus, variable amounts of BACE1 labeled dystrophic neurites were seen among sorfra/Amylo-Glo co-labeled plaques, while some Amylo-Glo heavily stained (likely burnout) plaques and pure sorfra plaques contained no or few BACE1 labeled elements (Supplemental Figure 20D, E, F).

Following the above microscopic characterization, we carried out correlative densitometry for sorfra and BACE1 labeling among the colocalized plaques using double immunofluorescently stained temporal lobe sections (Supplemental Figure 21A, B, C). Among the plaques quantified in CA1 and subiculum, there existed a negative correlation between normalized values (% of maximal density) of sorfra and BACE1 labeling among the plaques (P<0.0001, r=-0.24, n=495). Among the plaques quantified in the temporal neocortex, a trend of negative correlation existed between the variables but did not reach statistically significant difference (P=0.0862, r=-0.07, n=687) (Supplemental Figure 21D).

### Sorfra and pTau accumulation occurred with the loss of somatodendritic MAP2 labeling

Among the colocalized plaques in the neocortex, sorfra deposits often formed a rim surrounding the Aβ deposits (Figure 6B, D, F; Supplemental Figure 17). We speculated earlier that sorfra deposition might be associated with neurodegeneration especially of the somatodendritic compartment based on the normal expression pattern of sortilin (Hu et al., 2017; Xu et al., 2019; Shi et al., 2020). We addressed this possibility by examining sorfra and pTau accumulation relative to microtubule associated protein 2 (MAP2) expression in neuronal somata and dendrites. In sorfra/MAP2 double immunofluorescent preparation of sections with Amylo-Glo or DAPI counterstain (Figure 7A, Supplemental Figure 22A), there was an apparent loss of MAP2 labeled somata and dendritic profiles and DAPI stained nuclear profiles in the areas and with heavy loads of sorfra plaques in CA1, subiculum and amygdala (Figure 7A, A1; Supplemental Figure 22A1, A2) or in entorhinal cortical and neocortical regions (Figure 7A4, Supplemental Figure 22A5). At the sorfra plaque loci, MAP2 and DAPI labeling were absent or reduced (Figure 7A1, A2, A3, A4; Supplemental Figure 22A1, A2, A3, A5). At high magnifications, sortilin and MAP2 labeling were colocalized in neuronal somata and the dendritic tree in healthy-looking neurons in regions with no or few sorfra plaques (Figure 7A5; Supplemental Figure 22A3, A4). Sortilin/MAP2 double-labeled dendritic processes disappeared abruptly around the plaque border, or became apparently thinned as they passed across the plaques (Figure 7A1, A2, A4; Supplemental Figure 22A2, A3). Remnants of sortilin and MAP2 co-labeled processes could be observed inside some plaques (Figure 22A1, A5).

**Figure 7.**
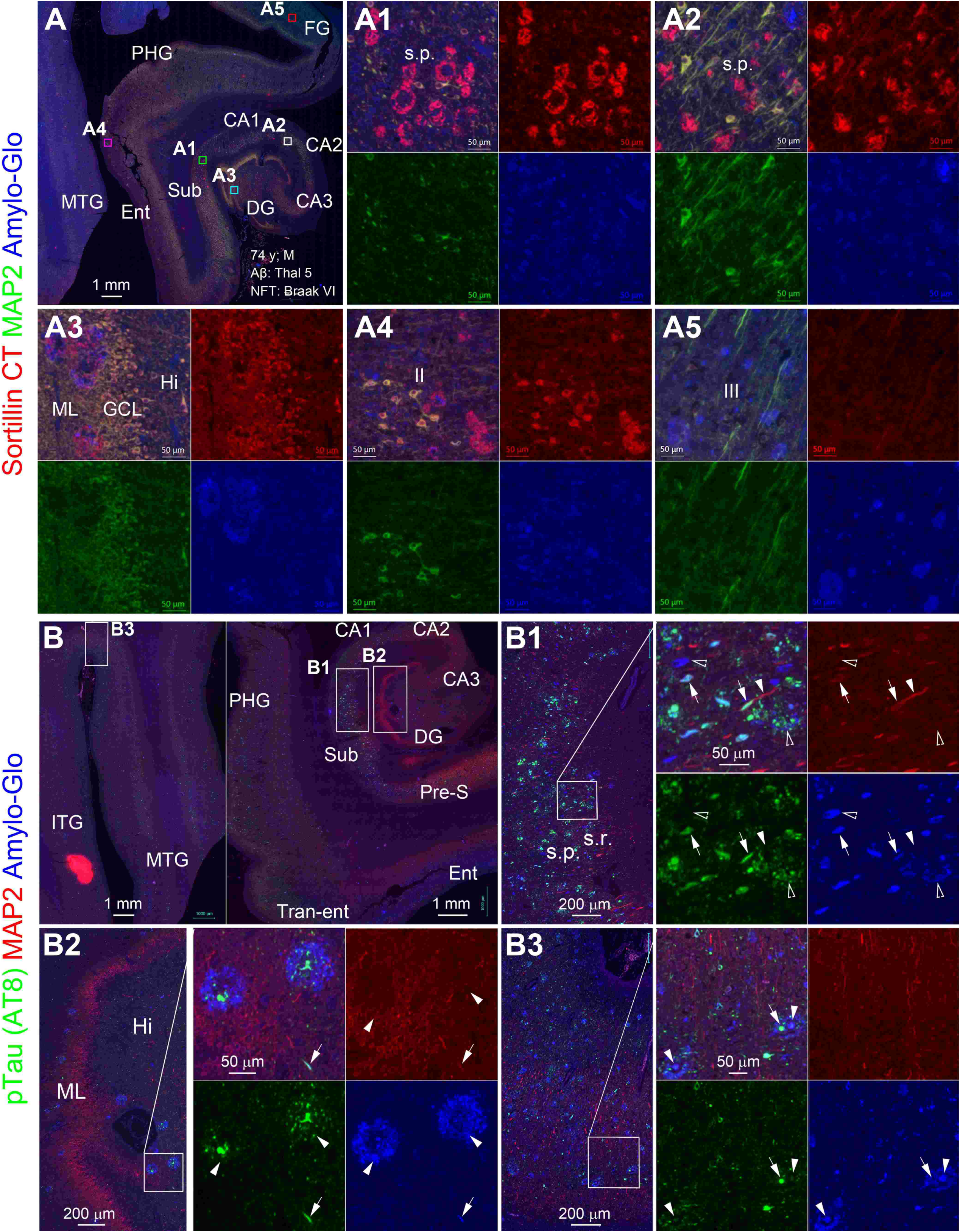
Sorfra and pTau/tangle pathologies relative neuronal somatodendritic alteration. In sorfra and microtubule-associated protein 2 (MAP2) double immunofluorescence with Amylo-Glo counterstain. Areas with heavy loads of sorfra plaques (**A**), or the loci of sorfra plaques (**A1, A2, A3, A4**), are depleted of MAP2 labeling. However, sortilin and MAP2 colocalized neuronal somata and dendritic processes are present in the unaffected areas (**A3, A5**). At high magnifications, the co-labeled dendritic processes disappear abruptly as they enter the plaque areas or become thinned as they pass the plaques (**A2, A4**). In pTau and MAP2 double immunofluorescence with Amylo-Glo counterstain (**B, B1, B2, B3**), triple-labeled neuronal somata and neuritic profiles (pointed by arrows) could be observed at high magnifications, which represent mature tangle-bearing structures. Neuronal somata and neuritic profiles are also seen being labeled only for pTau (pointed by arrowheads), which appear to represent pretangle profiles. In addition, somal and neuritic profiles are seen to exhibit Amylo-Glo labeling only (open arrows), likely representing ghost tangle profiles. The Amylo-Glo labeling appears to be “amorphic” among the plaque profiles in the dentate molecular layer (**A3, B2**) and in the entorhinal cortical and neocortical areas (**A4, A5, B2, B3**), which should be related to Aβ deposition, at least in part. Notably, pTau and MAP2 labeling can present segmentally in the same dendritic processes (**B1, B2**, pointed by arrows). Abbreviations: CA1-3: subregions of cornu Ammonis, DG: dentate gyrus, Hi: hilus; ML: molecular layer; GCL: granule cell layer, Pro-S: prosubiculum, Sub: subiculum, Pre-S: presubiculum, MTG: middle temporal gyrus, ITG: inferior temporal gyrus; Ent: entorhinal cortex; II-IV: cortical layers; Amyg: Amygdala; s.o.: stratum oriens, s.p.: stratum pyramidale; s.r.: stratum radiatum. Scale bars are indicated. Scale bars are as indicated.

In pTau/MAP2 double immunofluorescence with Amylo-Glo counterstain, tangle-like neuritic profiles labeled by pTau, Amylo-Glo or both occurred in clusters, with these profiles exhibited no or little MAP2 co-labeling in general (Figure 7B, B1, B2, B3; Supplemental 22B, B1). Specifically, MAP immunolabeling was not colocalized in the somal and neuritic profiles that were singly labeled by Amylo-Glo (Figure 7B1, B3), which represent ghost tangle profiles (Yang et al., 2024). A complementary pattern of pTau/MAP2 colocalization was observed in some neuronal somata and dendritic processes. In this case, the pTau labeling was concentrated in the soma of a neuron wherein the MAP2 labeling was depleted, whereas a mixed pTau/MAP2 labeling occurred along the dendritic trunk. In other cases, a segmental complementary pTau/MAP2 co-labeling was observed in dendritic processes that were not trackable to neuronal somata, including around the clusters of tangle-bearing neurites (Figure 7B, B1, B2; Supplemental Figure 22B1, B3). DAPI labeled nuclei appeared to be apparently lost in areas with heavy load of sorfra plaques (Supplemental Figure A1, A5, B1).

### Sorfra deposits occurred inside astrocytes but not microglia in peri-plaque areas

Finally, we explored potential glial clearance of sorfra deposits with double immunofluorescence. Sorfra deposits were observed inside some astrocytes around the plaques immunolabeled with glial fibrillary acidic protein (GFAP) antibody in paraffin sections from four brains (Figure 8A, A1, A2, B, B1, B2; Supplemental Figures 23A, A1, A2; 24A, A1, A2, B, B1, B2, B3). However, sorfra deposits were not observed in microglia immunolabeled with an Iba-1 antibody in the sections (Figure 8C, C1, C2, C3; Supplemental Figures 23B, B1, B2; 25A, A1, A2, A3, B, B1, B2).

**Figure 8.**
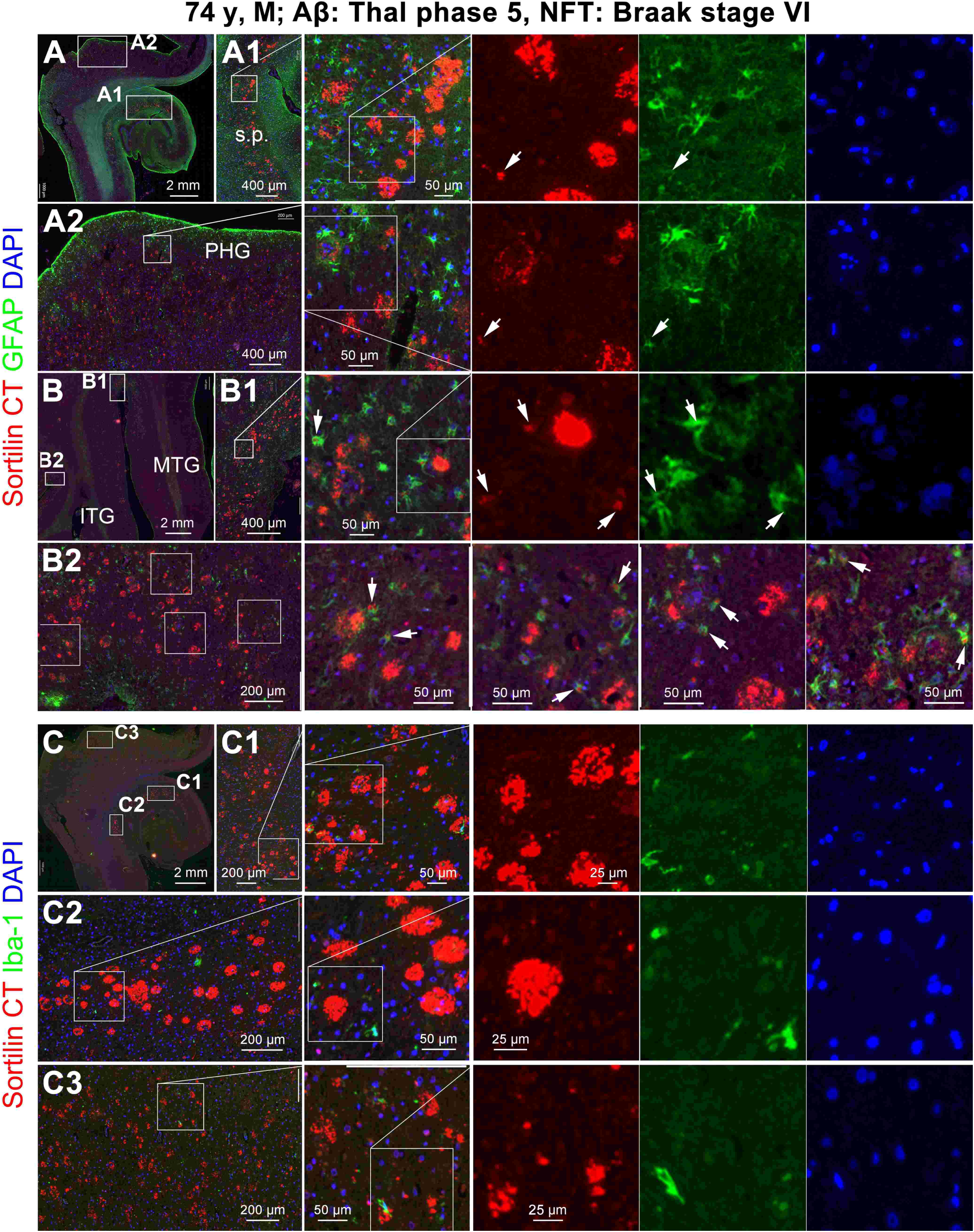
Double immunofluorescent characterization on potential glial phagocytosis of sorfra deposits. Representative images show double immunofluorescence for sorfra deposits and astrocytes labeled by the glial fibrillary acidic protein (GFAP) antibody (**A, A1, A2, B, B1, B2**, and enlarged panels as indicated), and sorfra deposits and microglia labeled by the Iba-1 antibody (**C, C1, C2, C3,** and enlarged views), with DAPI counterstain. Small chunks of sorfra deposits (pointed by arrows) occur inside some GFAP labeled astrocytes in the vicinity of the sorfra plaques in the hippocampal formation (**A1**) and temporal neocortical areas (**A2, B, B1, B2**). However, no sorfra deposits are detectable in Iba-1 labeled microglia in the hippocampal formation (**C, C1, C2**) or the temporal neocortex (**C, C3**). Abbreviations: PHG: parahippocampal gyrus, MTG: middle temporal gyrus, ITG: inferior temporal gyrus; s.p.: stratum pyramidale. Scale bars are as indicated.

## DISCUSSION

In neuroanatomical terms, neuritic plaques can be viewed as a lesion composed of abnormal neurites and extracellularly deposited amorphic material. Neurites are generally referred to as neuronal processes, with morphologically distorted axonal and dendritic neurites seen in many neuropathological conditions. Aβ plaques especially the so-called compact plaques have patch-like and size-limited Aβ deposition associated with dystrophic neurites as documented in human brain, rodent models of AD, and in the brains of many natural animals, which could be called neuritic plaques (Chambers et al., 2015; Fiock et al., 2020; Mesquita et al., 2021; Nakamura, 1996; Selkoe et al., 1987; Takahashi et al., 2022). The “neuritic” Aβ plaques characterized so far in humans, transgenic AD models and aged monkeys contain axonal dystrophic neurites expressing BACE1 (Zhang et al., 2009; Cai et al., 2010; Cai et al., 2012; Sadleir et al., 2016). The extent to which Aβ plaques represent or transform into the traditional silver-stained neuritic plaques remains a fundamental pathogenic question in AD, since the latter are highly implicated for pathological diagnosis and cognitive impairment (Hyman et al., 2012; Montine et al., 2012; Nelson et al., 2009; Tsering & Prokop, 2024). Braak and colleagues compared amyloid plaques and neuritic plaques in the human brain between consecutive paraffin sections and pointed out that most of the former type lacked silver stained dystrophic neurites (Braak et al., 1989).

Braak staging through cross-sectional analysis of pTau and silver stained tangle pathology in human brains relative to age and disease status established a tight link of neuronal tauopathy to tangle formation and neuritic plaque development (Braak et al., 2006; Braak & Del Tredici, 2011). Notably, fully “mature” neuritic plaques appear to be only prominent in humans. Animals including nonhuman primates can develop substantial cerebral Aβ deposition and some neuronal tauopathy with age, but rarely recapitulate the forms and extent of tangle and neuritic plaque lesions seen in AD human brains (Barnes et al., 2024; Ferrer, 2024). Besides the hierarchic progression from the limbic to and across the neocortical regions, a sulcal valley to gyral hilltop differential distribution of cortical neuritic plaques was noticed at least by the early 1960s (Beach, 2022; McMenemy, 1963). The concept of neuritic plaque and tangle evolution was proposed since the time these lesions were discovered (Fiala, 2007; Walker, 2020; Beach, 2022). Thus, neuritic plaques were thought to develop from primitive to mature then burnout stages, while tangle pathogenesis proceeds through pretangle, mature tangle and ghost tangle stages (Stokin et al., 2005; Terry & Wiśniewski, 1970; Munoz & Wang, 1992; Gómez-Ramos & Morán, 1998; Moloney et al., 2021).

The present study used two sortilin C-terminal antibodies to validate the sorfra pathology identified earlier by our group (Hu et al., 2017). These two antibodies showed essentially identical performance in immunoblot and immunohistochemistry. Thus, they immunoblotted sortilin fragments migrated at ∼15 kDa and ∼55 kDa, as well as the ∼100 kDa full-length holoprotein, and labeled extracellular deposits arranged as small and isolated fibrils to microscopically prominent plaques (mostly 50-100 μm and rarely exceeding 200 μm in diameter). They also marked intraneuronal inclusions, with some resembling the granulovacuolar degeneration bodies(Jiang et al., 2022). In addition, neuronal somata and dendrites were labeled in brains or brain areas with less or no extracellular sorfra pathology, due to a cross-immunoreaction with the full-length sortilin protein. Because the antibody labeling can be absorbed by the antigenic peptide encompassing the 800-830 a.a. sequence of the C-terminal, the extracellular sorfra deposits are likely formed by the smaller, i.e., the 15 kDa, fragment (Hu et al., 2017). The holoprotein cross-reactivity is a pitfall of these antibodies in a sense that it limits their utility for the analysis of sorfra in biofluids using methods such as enzyme-linked immunosorbnent or immunodot assay. On the other hand, this cross-reactivity allows an appraisal of the cellular origin of sorfra.

Consistent with earlier mapping data obtained from cryostat sections (Tu et al., 2020), a parallel regional progression was observed for sorfra plaques, pTau immunolabeling, silver-stained and tangle tracer stained somal and neuritic lesions. Such a parallelism was seen across a single temporal lobe section covering the hippocampal formation, entorhinal cortex and temporal neocortex, or an occipital lobe section covering the primary and secondary visual cortices. A transitional pattern between sulcal valley to gyral hilltop regions was impressive and seen in many local neocortical areas in the brains with NFT pathology from Braak stages III-VI. Quantitatively, the burdens of sorfra and pTau were positively correlated among cortical zones; and there existed a parallel decline in the burden of sorfra plaques and Gallyas silver stained profiles, but not that of Aβ plaques, in cortical areas as moving from the sulcal valley to gyral hilltop. By cross-examination of consecutive paraffin sections, sorfra plaques were found to closely match with silver-stained neuritic plaques in location, size and shape, especially evident in the plaques in the limbic structures that contained thick tangle-bearing neurites. Most typical compact Aβ plaques in the neocortex contained few or no tangle-bearing neurites or neurofibrillary threads, consistent with the observation described by Braak and colleagues (Braak et al., 1989). Further, sorfra plaques were associated with tangle-bearing dystrophic neurites and neuritic threads in pTau immunofluorescence with the counterstain of a selective tangle tracer TNIR7-1A. Taken together, sorfra plaque development is related to neuronal tangle pathogenesis. To explore sorfra deposition relative to the transformation of Aβ plaques into neuritic plaques, we first assessed Aβ/sorfra immunolabeling with Amylo-Glo stain in sections with and without formic acid treatment. The results indicated that sorfra deposits are nonamyloid in nature. Along with further assessment of sorfra immunolabeling with the staining of a selective amyloid tracer DANIR 8c, it was fairly clear that sorfra and Aβ can form colocalized as well as independent plaques. To overcome the concern that microscopic section study may “miss the forest for the trees”, tissue clearance three-dimensional imaging technology was used, which confirmed the above-mentioned differential plaque forming pattern in AD neocortex. Densitometric analyses established a negative correlation of sorfra deposition relative to Aβ deposition and amyloid tracer staining among the colocalized plaques. Given that Aβ plaques develop earlier than tau/tangle pathology in the neocortex and sorfra deposition is related to the latter as mentioned above, the transformation of amyloid plaques to neuritic plaques appears to proceed by a decrease of the amyloid Aβ, but an increase of the nonamyloid sorfra, components. The mechanism underlying the former trend remains to be investigated further. However, this could be due to a reduced Aβ production, given that a trend of inverse correlation was also found between sorfra deposition and BACE1 labeling of dystrophic presynaptic terminals (Cai et al., 2010; Cai et al., 2012; Ferrer, 2023; Jordà-Siquier et al., 2022; Sadleir et al., 2016; Zhang et al., 2009).

The morphology of sorfra plaques showed a certain extent of difference between the neocortex and the hippocampal formation and amygdala (limbic structures). As such, in double labeling preparation, sorfra deposits formed a rim surrounding the Aβ or amyloid tracer labeling among many colocalized plaques in the neocortex. Hippocampal and amygdalar sorfra plaques were often massed in mix with Aβ or amyloid tracer labeling when the latter was present. Also as seen in double immunofluorescence, sorfra deposition occurred along with a loss of MAP2 labeled neuronal somata and dendritic processes, regionally or locally depending on the load of sorfra plaques, resulting in a complementary pattern of MAP2 and sorfra labeling across the limbic and neocortical regions recognizable at low magnifications. The cellular origin of sorfra from degenerating somatodendritic components could be perceived by the loss and thinning of MAP2/sortilin co-labeled dendritic processes around individual plaques. This perception was further supported by the assessment of MAP2 relative to pTau and tangle tracer labeling. Thus, MAP2 labeling was reduced in neuronal somata and dendritic processes along with pTau accumulation and tangle formation. The finding that sorfra deposits are phagocytized by astrocytes is in line with a neurodegenerative event that triggers cellular clearance response. Since astrocytic processes are a part of synaptic trisome, these glial cells may be activated in response to tangle-induced somatodendritic degeneration. Taken together, the formation of a sorfra rim among the cortical Aβ plaques could be explained as a result of pTau/tangle pathogenesis in the remaining neurons surrounding the plaques. The earlier and severer tauopathy in the limbic structures could underlie a greater sorfra formation and deposition as massed plaques in these regions. Further, with the spreading and worsening of tau and tangle pathology, sorfra or neuritic plaques could develop independent of Aβ plaques in the human brain.

## CONCLUSION

The present study determined sorfra pathogenesis relative to tau and amyloid in the human brains with late stages AD pathologies using multiple combinations of labeling methods and comparative image and quantitative analyses. Sorfra formation and deposition proceed regionally and locally in parallel with pTau and tangle pathogenesis and are cellularly linked to neurodegeneration, especially the degradation of the somatodendritic components. As such, sorfra deposition contributes to a nonamyloid growth of the Aβ plaques and also forms nonamyloid neuritic plaques. These findings extend new insights into the integration of Aβ and pTau pathologies and the evolution of neuritic plaques in Alzheimer’s disease.

## DATA AVAILABILITY STATEMENT

The raw data supporting the conclusions of this article will be made available by the authors, without undue reservation. Extended data presented as Supplemental figures 1-25 is provided online.

## ETHICS STATEMENT

Written consent for whole body donation for medical education and research was obtained from the donors or next of kin of subjects in compliance with the body/organ donation laws and regulations set by Chinese government. Use of postmortem human brains was approved by the Ethics Committee for Research and Education at Xiangya School of Medicine, in compliance with the Code of Ethics of the World Medical Association (Declaration of Helsinki).

## AUTHOR CONTRIBUTIONS

All authors read and approved the final manuscript. Conceptualization: XXY; Methodology: QLZ, YW, SC, ZPS, XLC, TT; Formal analysis and investigation: QLZ, TT, YL; Writing - original draft preparation: QLZ; Writing - review and editing: XXY, JM; Funding acquisition: XXY, YZ, AP; Resources: YZ, MC, JW.

## ACKNOWLEDGMENTS

We thank Xiao-Hua Tang for help with Motic light microscopic imaging, and Dan-Dan Hu, Qiang Li, Qian-Li Shen for human brain banking.

