## supplemental figures for "Sortilin C-terminal fragment deposition depicts tangle-related nonamyloid neuritic plaque growth in Alzheimer’s disease"

### Supplemental Figures 1-25

#### Sortilin C-terminal fragment deposition depicts tangle-related nonamyloid neuritic plaque growth in Alzheimer's disease

Qi-Lei Zhang[1], Yan Wang[1], Sidiki Coulibaly[1], Zhong-Ping Sun[1], Xiao-Lu Cai[1], Tian Tu[2], Aihua Pan[1], Meng-Chao Cui[3], Jim Manavis[4], Jian Wang[5], Yang Zhang[6], Xiao-Pin Wang[6] and Xiao-Xin Yan[1, 5, 6*]

[1]Department of Anatomy and Neurobiology, Central South University Xiangya School of Medicine, Changsha, Hunan 410013, China
[2]Department of Neurology, Central South University Xiangya Hospital, Changsha, Hunan 410008, China
[3]Key Laboratory of Radiopharmaceuticals, Ministry of Education, College of Chemistry, Beijing Normal University, Beijing 100875, China
[4]Faculty of Health and Medical Sciences, The University of Adelaide, Adelaide, SA 5005, Australia
[5]Department of Pathology, Hunan Guangxiu Hospital, Changsha, Hunan 410119, China
[6]Department of Psychiatry, National Clinical Research Center for Mental Disorders, and Brain Bank for Psychiatric Diseases, The Second Xiangya Hospital of Central South University, Changsha Hunan 410011, China

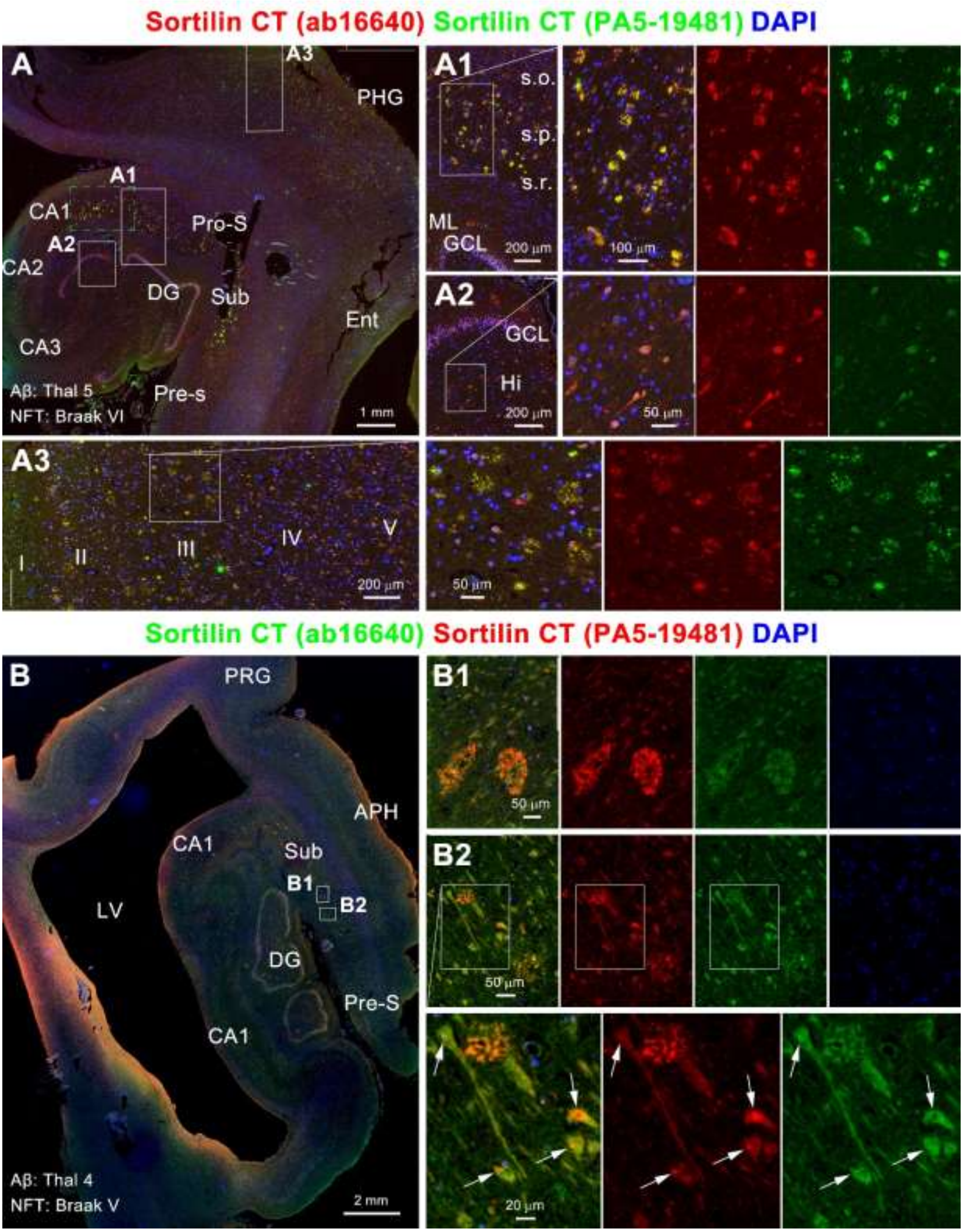


**Supplemental Figure 1: Double immunofluorescent characterization of the two sortilin C-terminal antibodies with the tyramide signal amplification (TSA) method in paraffin sections from two brains.** The immunolabeling is alternatively visualized using two polymer-horseradish peroxidase (HRP) conjugated secondary antibodies as indicated, which displays identical labeling of sorfra plaques, neuronal somata and dendrites, and intracellular inclusions (pointed by arrows). Abbreviations are as defined in main Figures 1 and 2.

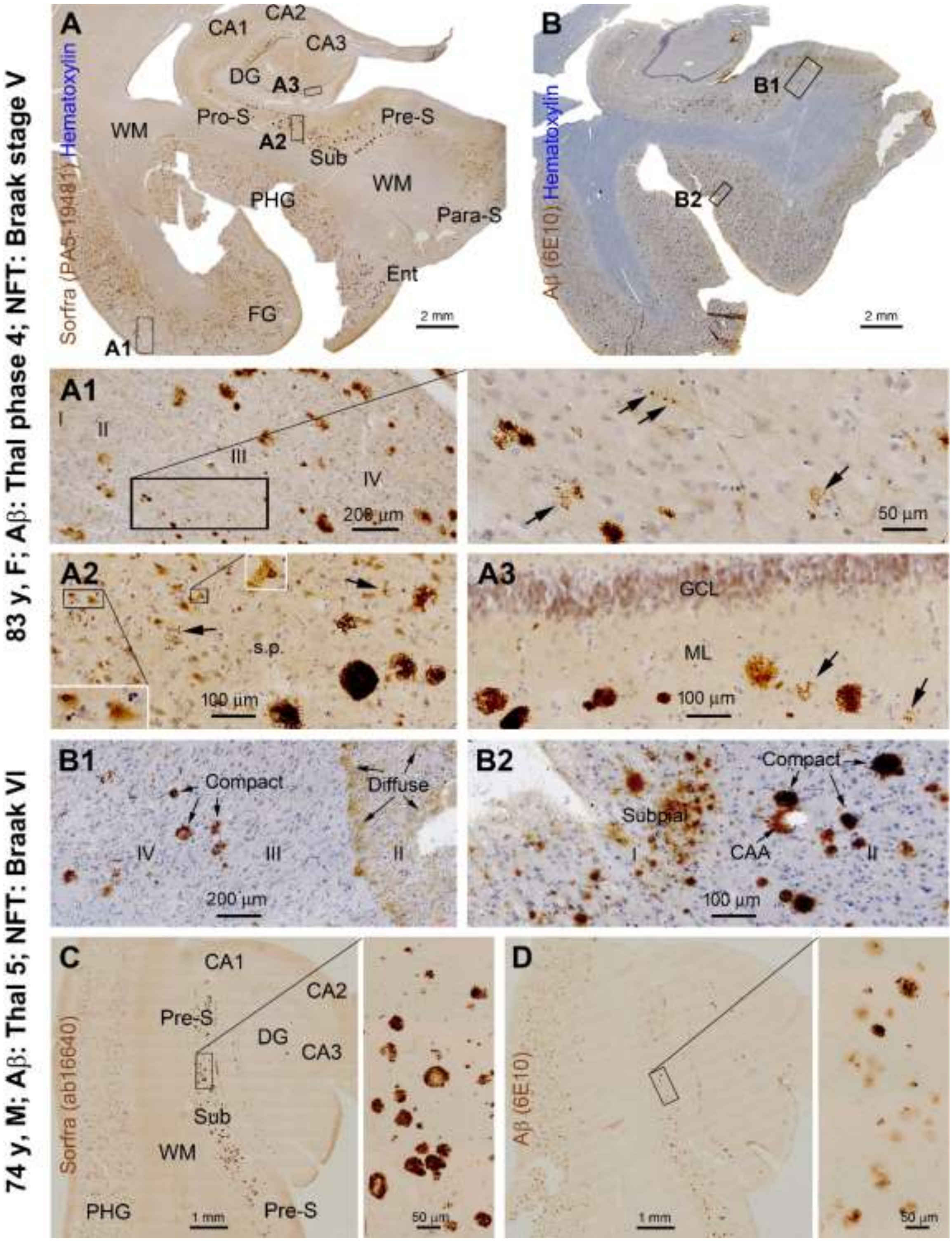


**Supplemental Figure 2: Deposition of sortilin C-terminal fragments (sorfra) and β-amyloid (Aβ) in cryostat temporal lobe sections from two AD brains.** Sorfra pathology is assessed with two sortilin antibodies (**A, C**), with Aβ labeled by the 6E10 antibody (**B, B1, B2, D**). Sorfra and Aβ plaques are present across the neocortex, entorhinal cortex and hippocampal subregions. Sorfra deposits arrange as discrete fibrillary profiles to massed plaques (**A1, A2, A3, C1**, arrows). Intracellular inclusions are seen in some neurons (**A2**, inserts). Aβ pathology includes compact, diffuse and subpial plaques, and cerebral amyloid angiopathy (CAA) (**B, B1, B2**). In the subiculum, sorfra plaques are more densely stained relative to Aβ plaques (**C, D**). Abbreviations are as defined in main Figures 1 and 2.

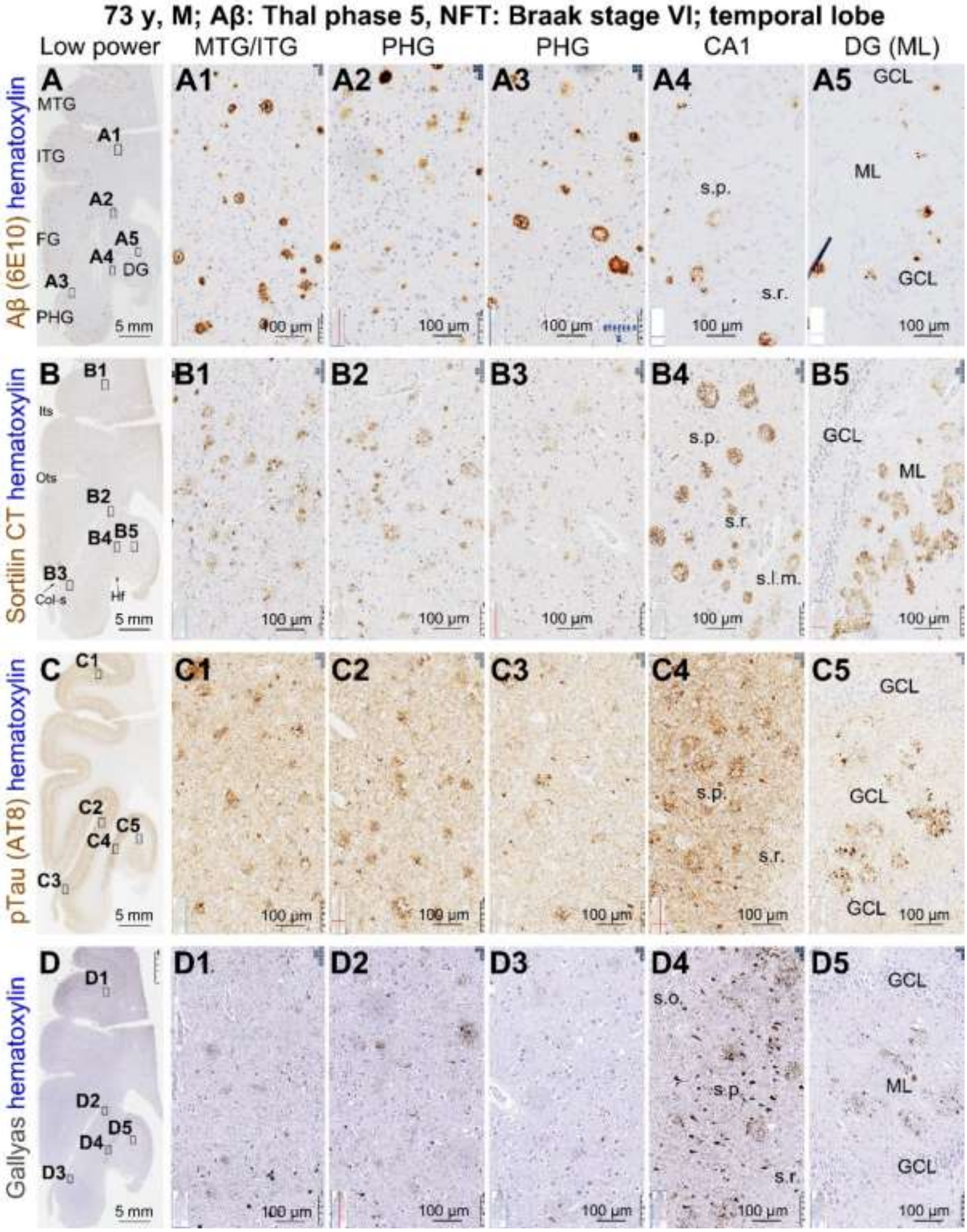


**Supplemental Figure 3: Distribution of Aβ, sorfra, pTau and silver tangle labeling in temporal lobe sections from an end-stage AD case.** Lesions are seen throughout the temporal neocortical gyri besides entorhinal and hippocampal areas (**A, B, C, D**). Cortical Aβ plaques appear more heavily labeled relative to their hippocampal counterpart (**A1-A5**), with an opposite trend seen in sorfra (**B1-B5**), pTau (**C1-C5**) and Gallyas (**D1-D5**) preparations. A sulcal valley to gyral hilltop difference in the amount of labeled profile along the collateral sulcus (Col-s) is not seen in Aβ (**A2, A**), but seen in sorfra (**B2, B3**), pTau (**C2, C3**) and Gallyas silver (**D2, D3**), preparations.

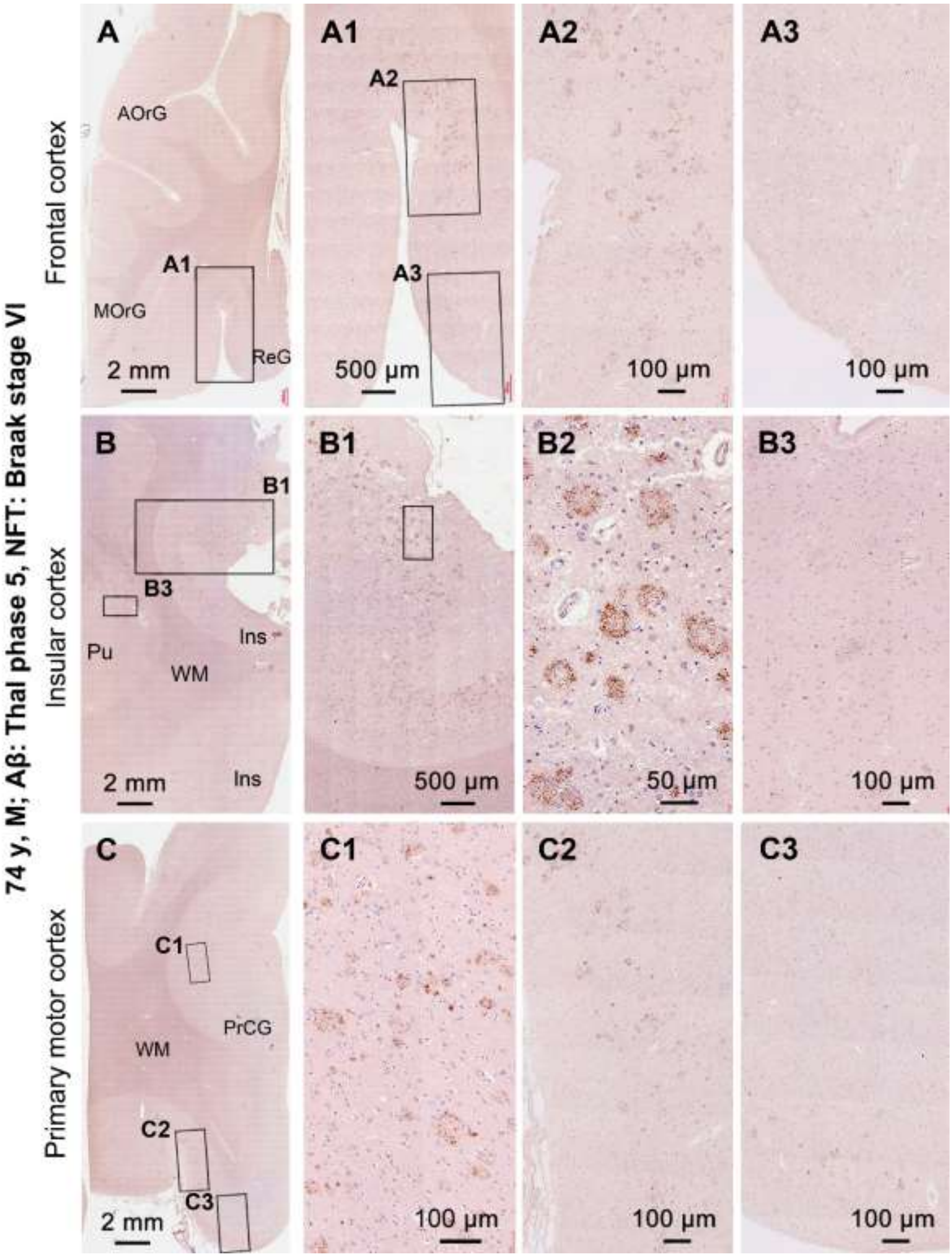


**Supplemental Figure 4: Distribution of sorfra plaques in cerebral neocortical regions in an end-stage AD case.** Sorfra plaques are observed in the frontal (**A**), insular (Ins) (**B**) and motor (**C**) cortical regions. The brain has a high score of tauopathy (Braak stage VI). Sorfra plaques are found in all the above neocortical regions. In each gyrus, the plaques are most densely distributed in the cortex surrounding the sulcal bottom (**A1, A2, B1, B2, C1, C2**). As moving from here towards the gyral hilltop, the density of the plaques reduces progressively (**A3, B3, C3**). AOrG: anterior orbit gyrus, MOrG: medical orbit gyrus, ReG: rectus gyrus, PrCG: precentral gyrus. Pu: putamen

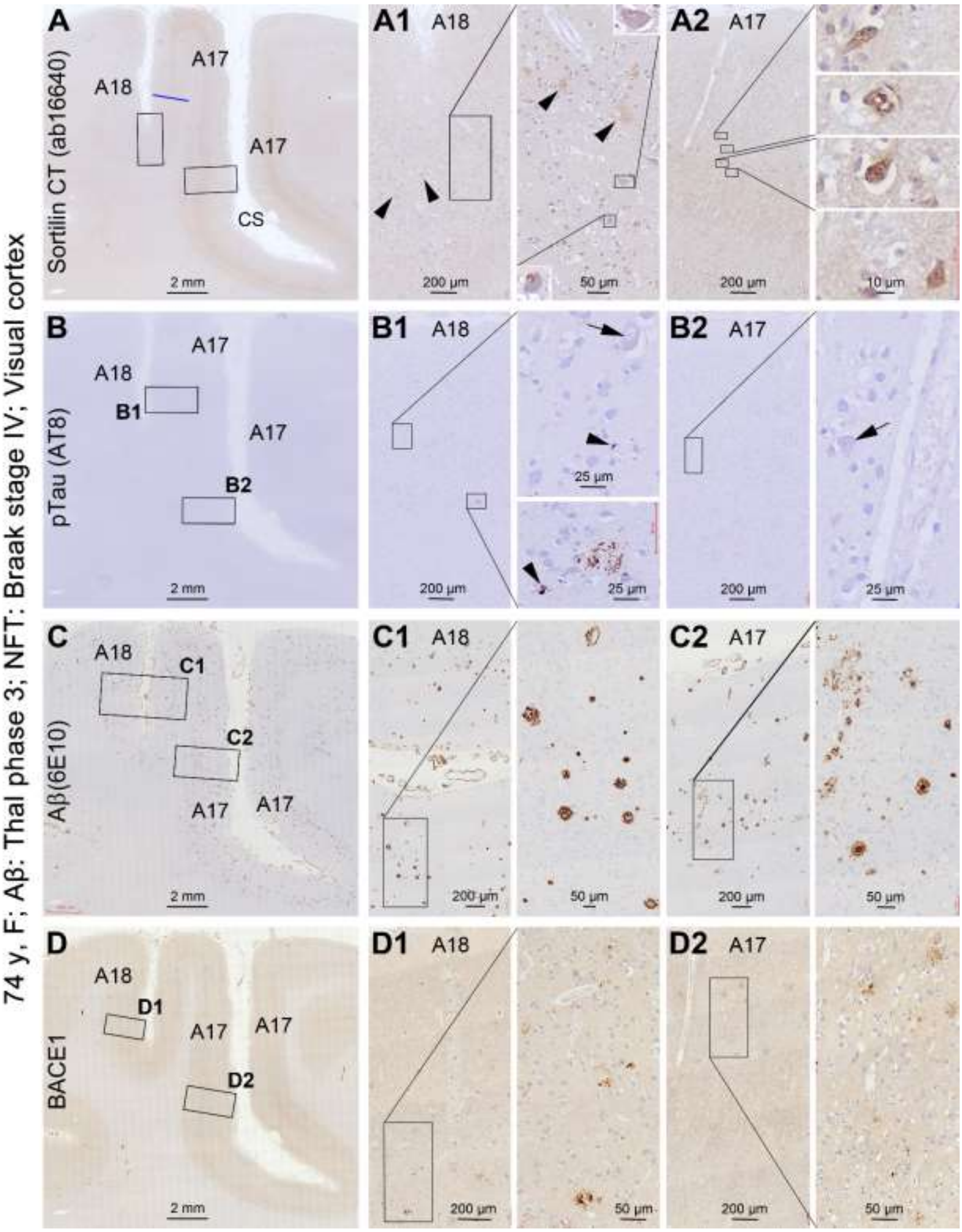


**Supplemental Figure 5: Differential distribution of sorfra and pTau labeling relative to Aβ and β-secretase 1 (BACE1) between areas 17 and 18 in a human brain with Thal phase 3 Aβ and Braak stage IV tau pathologies.** Sorfra plaques and pTau labeled neuritic clusters are seen in low number in area 18 (**A, A1**, arrowheads), but are not found in area 17 (**A, A2**). Granulovacuolar degeneration bodies are seen in both area 17 and 18 (**A1, A2**, inserts, arrows). Aβ plaques (**C, C1, C2**) and BACE1 labeled neuritic clusters (**D, D1, D2**) are concurrently observed in areas 17 and 18.

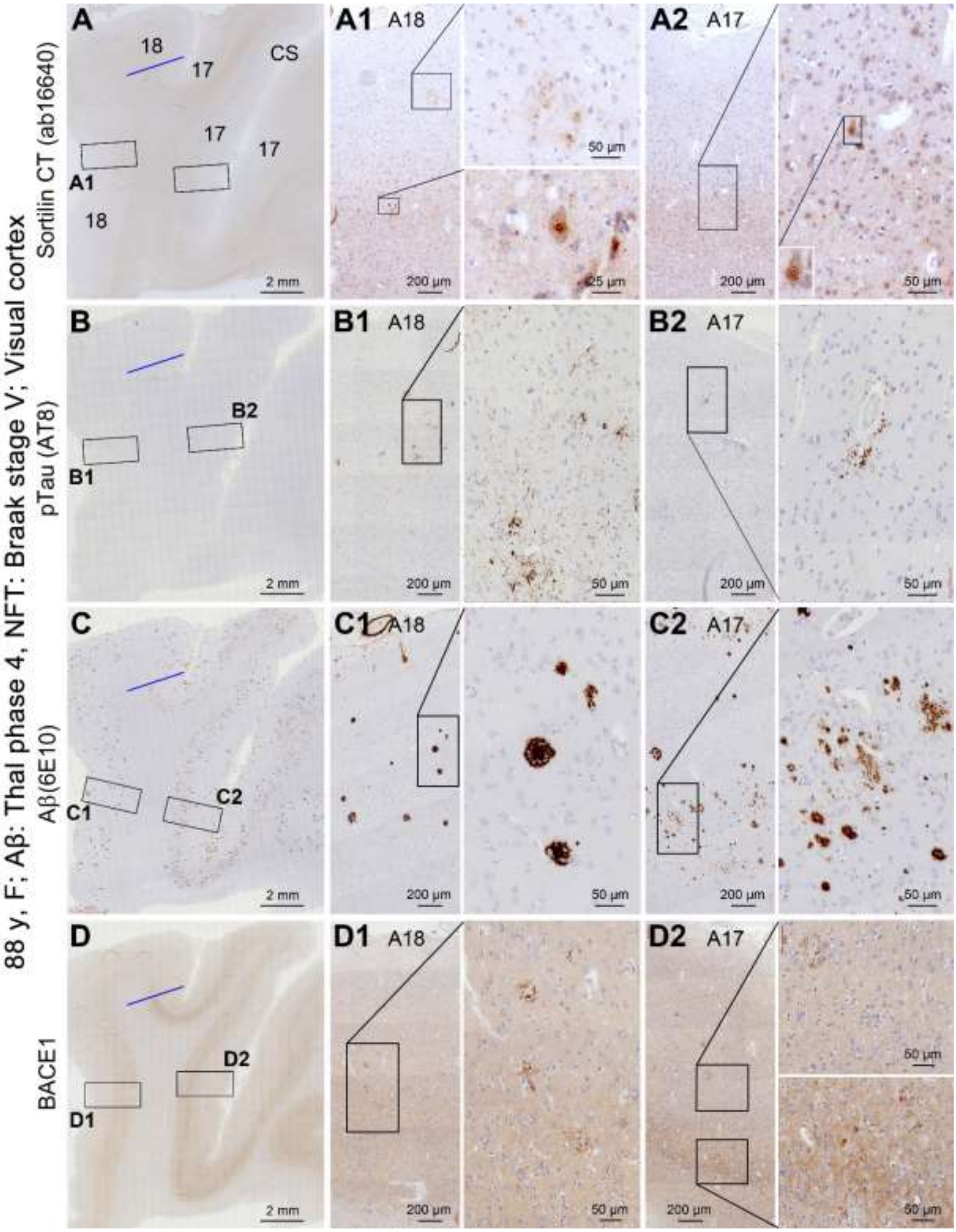


**Supplemental Figure 6: Differential distribution of sorfra and pTau labeling relative to Aβ and β-secretase 1 (BACE1) between areas 17 and 18 in a human brain with Thal phase 4 Aβ and Braak stage V tau pathologies.** Sorfra plaques and pTau labeled neuritic clusters are seen in both visual cortices, but noticeable dense in area 18 (**A, A1, B, B1**) than area 17 (**A, A2, B, B2**). Granulovacuolar degeneration bodies are seen in both visual cortical areas (inserts). Aβ plaques (**C, C1, C2**) and BACE1 labeled neuritic clusters (**D, D1, D2**) are present in areas 17 and 18.

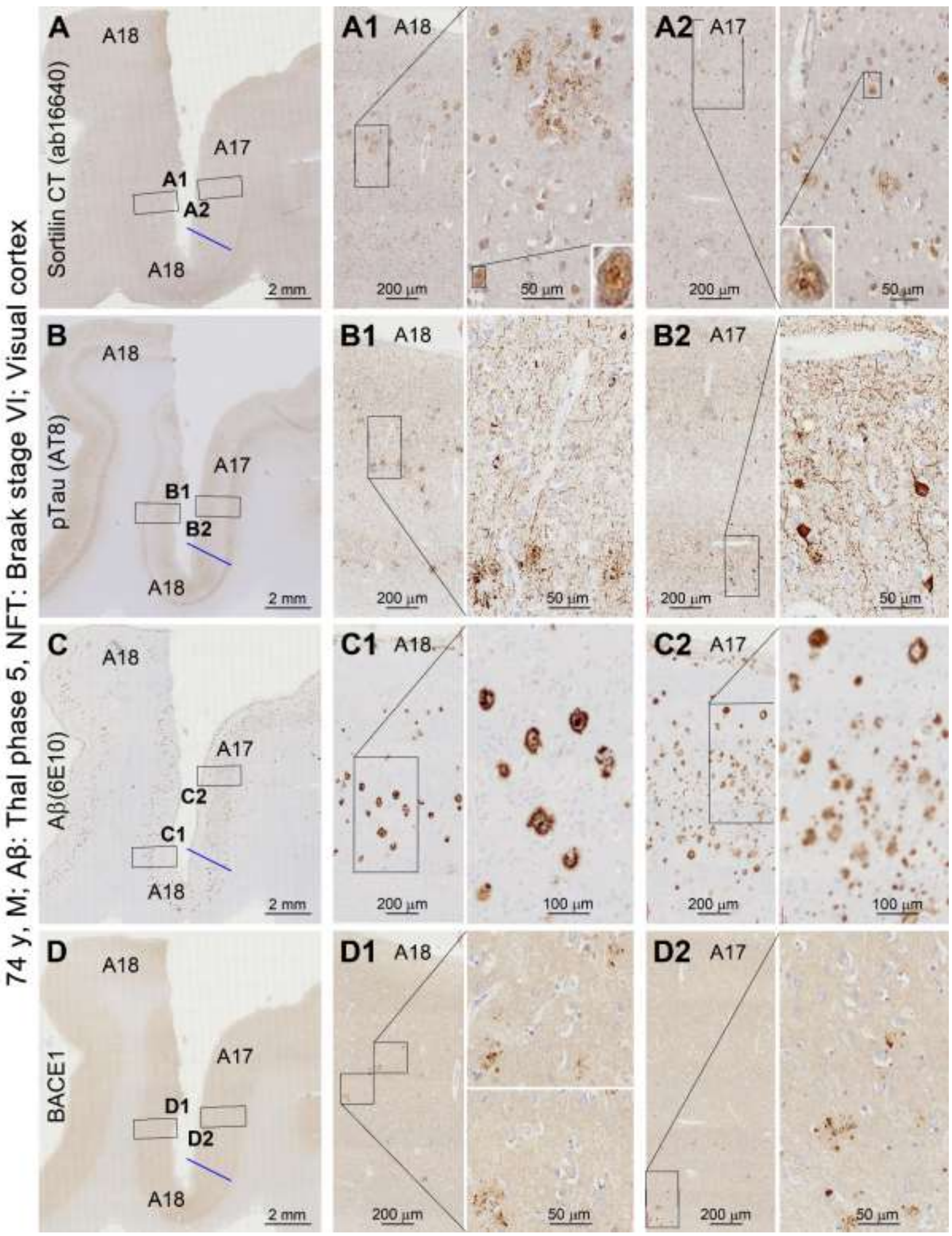


**Supplemental Figure 7: Sorfra, pTau, Aβ and β-secretase 1 (BACE1) labeling in areas 17 and 18 in a human brain with Thal phase 5 Aβ and Braak stage VI tau pathologies.** All four types of pathological profiles are present in areas 17 and 18 (**A, B, C, D**). The amount of sorfra and pTau labeling appears to be greater in area 18 than in area 17 (**A, A1, A2, B, B1, B2**). Aβ plaques appear largely as compact plaques in area 18 (**C, C1**), while those in area 17 appear mostly the diffuse type (**C, C2**). BACE1 labeled dystrophic neurites occur in clusters varying in size (**D, D1, C2**).

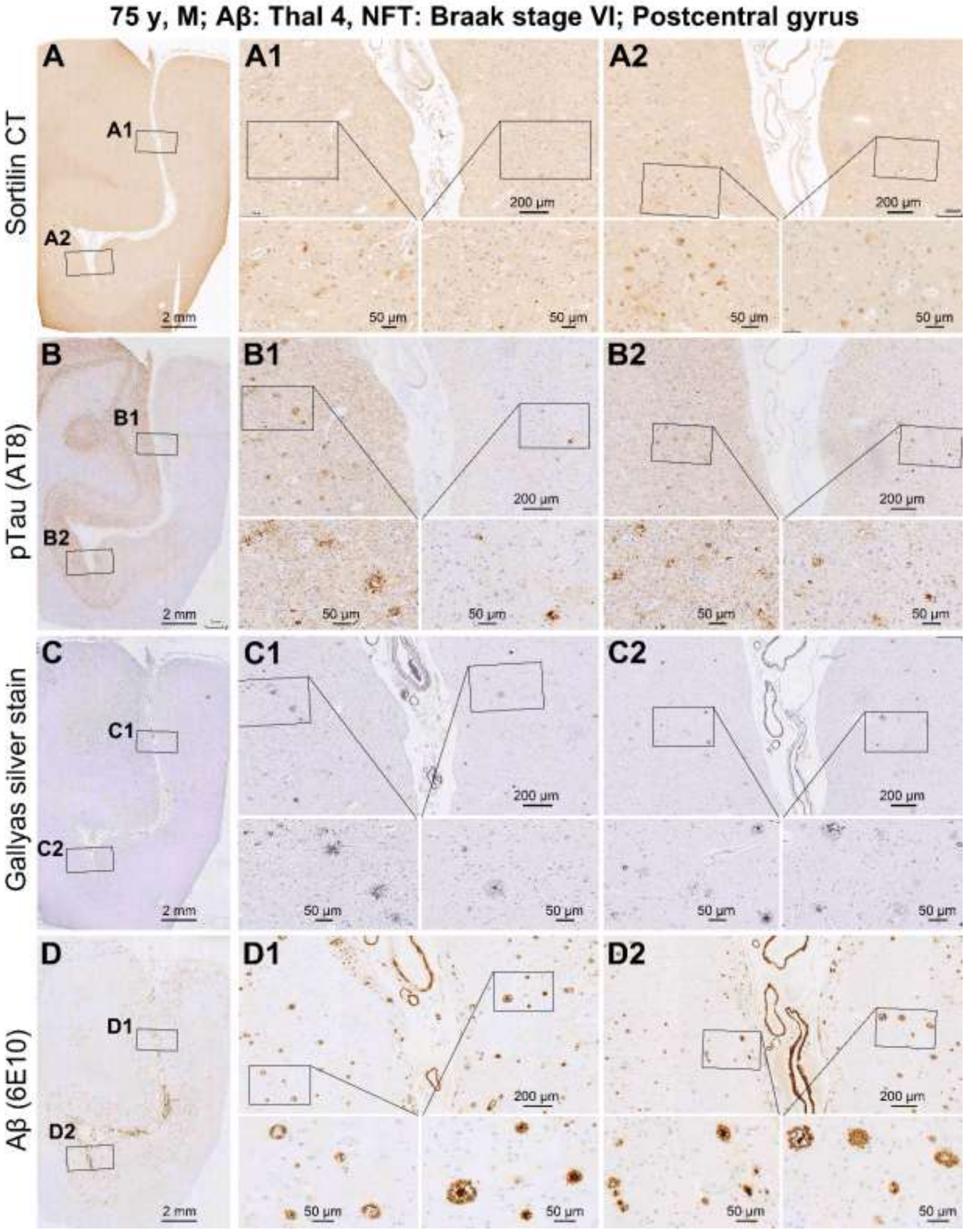


**Supplemental Figure 8: Differential distribution of sorfra, pTau and silver tangle labeling, but not Aβ labeling, between adjacent neocortical gyri.** Sections are from a brain with Braak stage VI and Thal phase 4 lesions. The extent of sorfra (**A, A1, A2**), pTau (**B, B1, B2**) and Gallyas silver (**C, C1, C2**) labeling appears greater in one side of the cortex relative the other bordering a cortical sulcus in the postcentral gyrus. The amount of Aβ plaques appear comparable between the two sides (**D, D1, D2**).

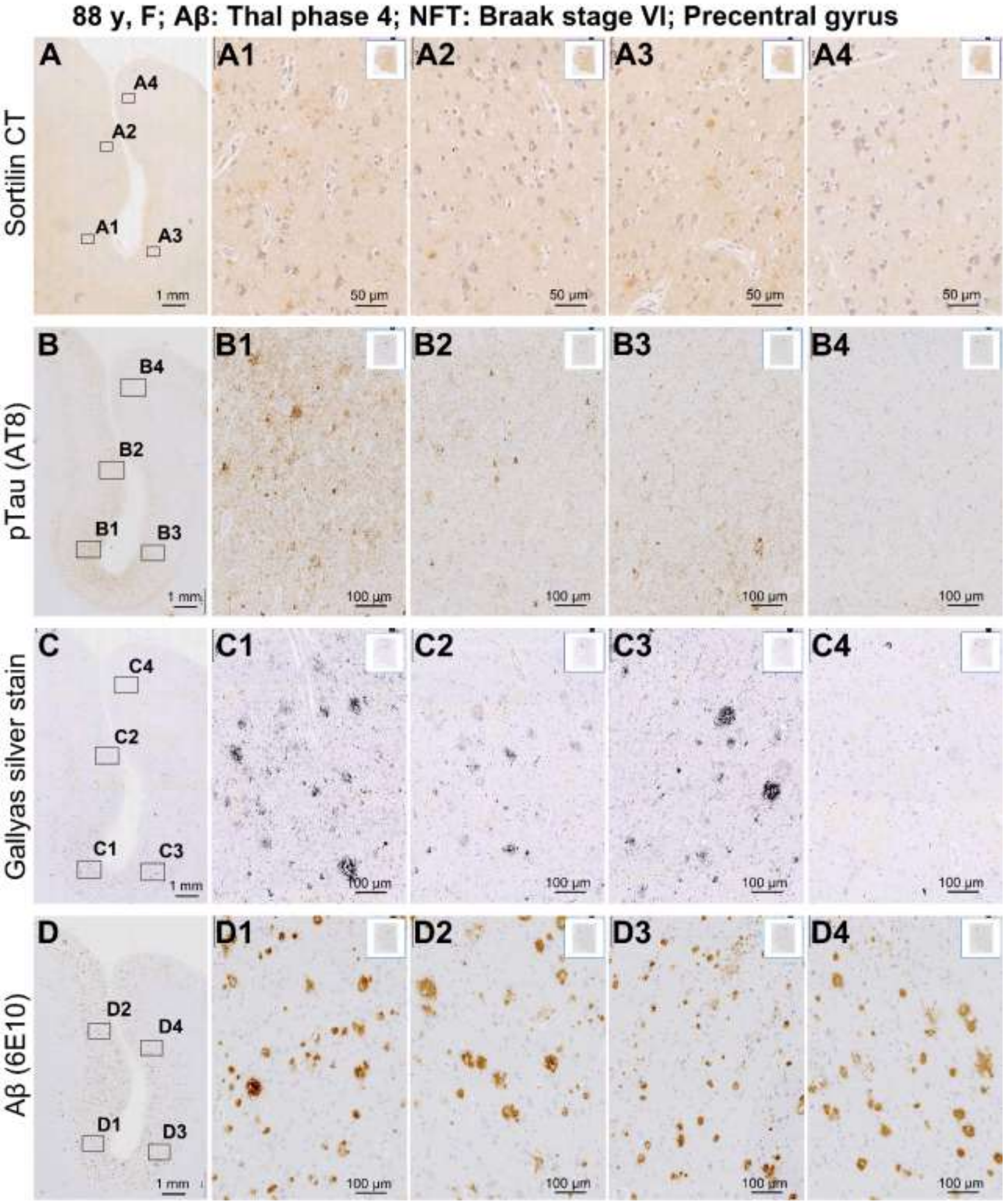


**Supplemental Figure 9: Further example of differential distribution of sorfra, pTau and silver tangle labeling, but not Aβ labeling, in local neocortical areas.** Sections are from a brain also with Braak stage VI and Thal phase 4 lesions. By comparing the image panels as indicated, there exists a differential distribution pattern in sorfra (**A, A1-5**), pTau (**B, B1-4**) and Gallyas silver (**C, C1-4**) labeling between neighboring gyri, and between the sulcal valley and gyral hilltop. This differential local distribution pattern is not evident in Aβ labeling (**D, D1-D4**).

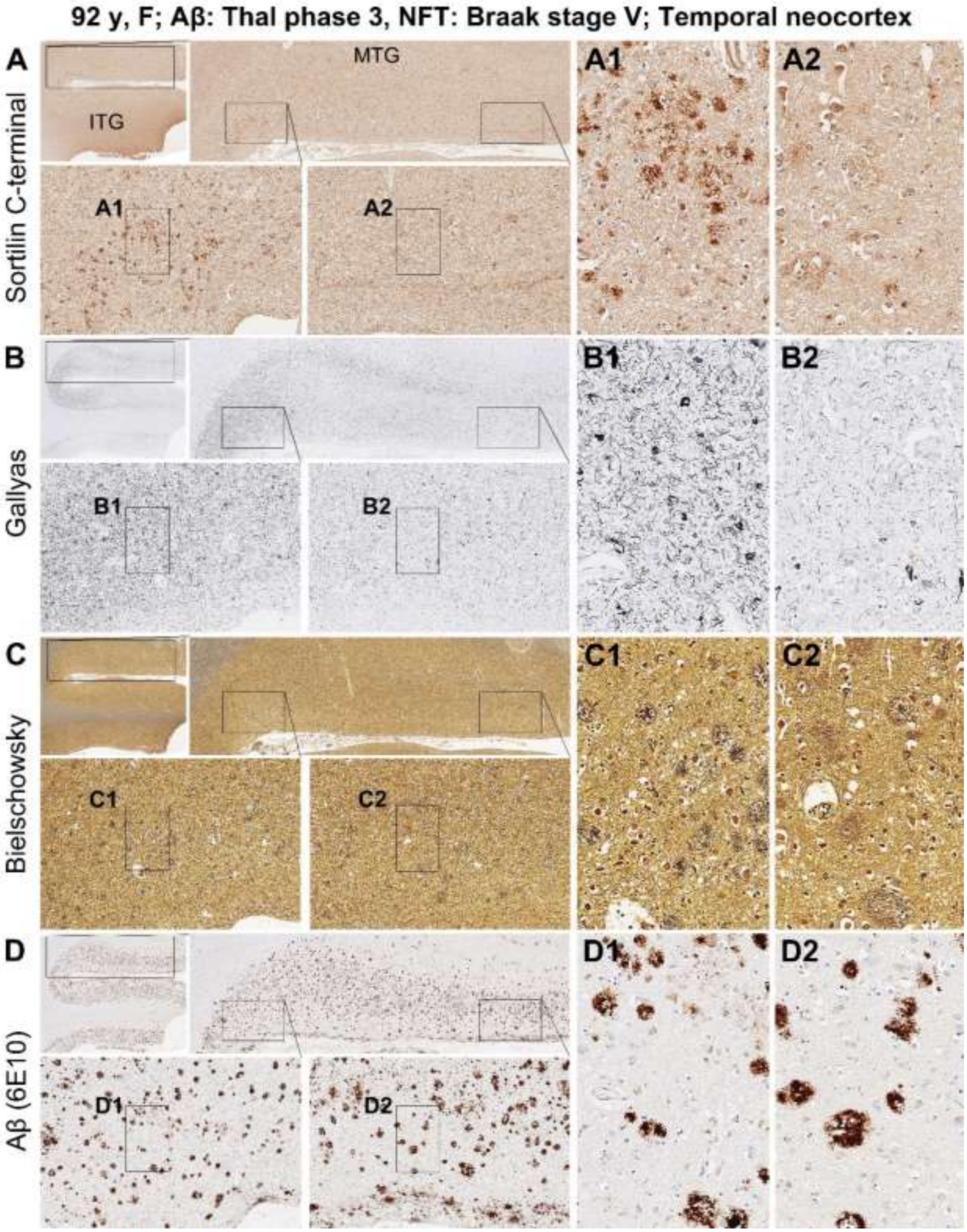


**Supplemental Figure 10: Differential distribution of sorfra plaques and neuritic plaques stained with Gallyas and Bielschowsky methods, but not Aβ plaques, along sulcal valley to gyral hilltop transition.** Braak stage and Thal phase of the brain are as indicated. Sorfra plaques, silver stained tangles and neuritic plaques are heavily present around the sulcal valley (**A, A1, B, B1, C, C1**), but remarkably reduced near the gyral hilltop (**A, A2, B, B2, C, C2**). The overall density of Aβ plaques appear to be comparable as moving from the valley to hilltop (**D, D1, D2**).

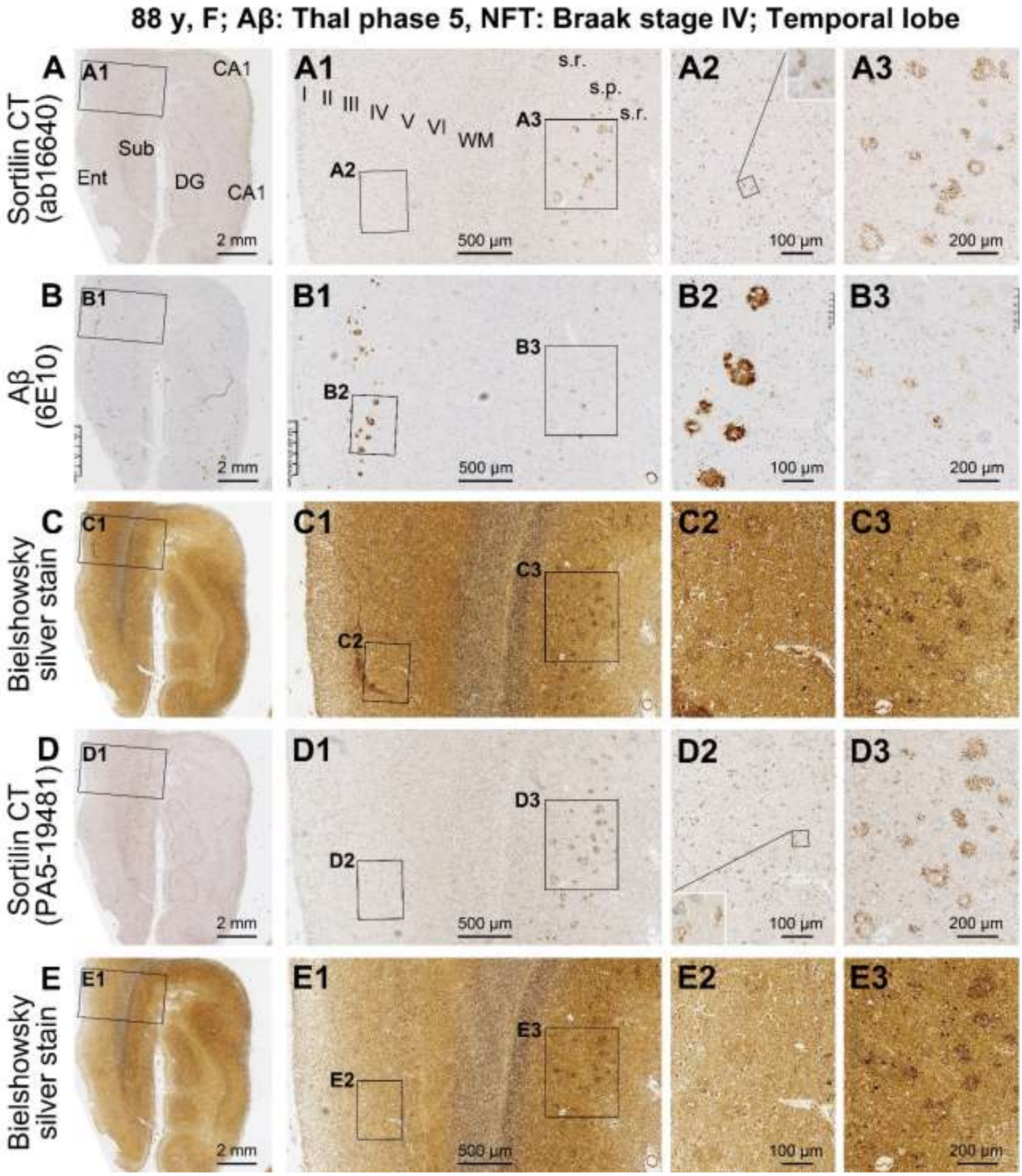


**Supplemental Figure 11: Assessing the anatomical matchiness of sorfra plaques, neuritic plaques and Aβ plaques using serial consecutive temporal lobe paraffin sections from a human brain with late stages of AD pathology.** In the temporal neocortex, heavily labeled Aβ plaques can occur in the absence of sorfra co-deposition and silver stained neuritic processes (**A, A1, A2, B, B1, B2, C, C1, C2, D, D1, D2, E, E1, E2**), although discrete sorfra deposition are present locally (**A2, D2,** inserts). In the subiculum and CA1 areas, sorfra plaques and silver stained neuritic plaques are well developed and mostly matchable in location, size and shape, whereas these plaques exhibit weak Aβ immunolabeling (**A3, B3, C3, D3, E3**).

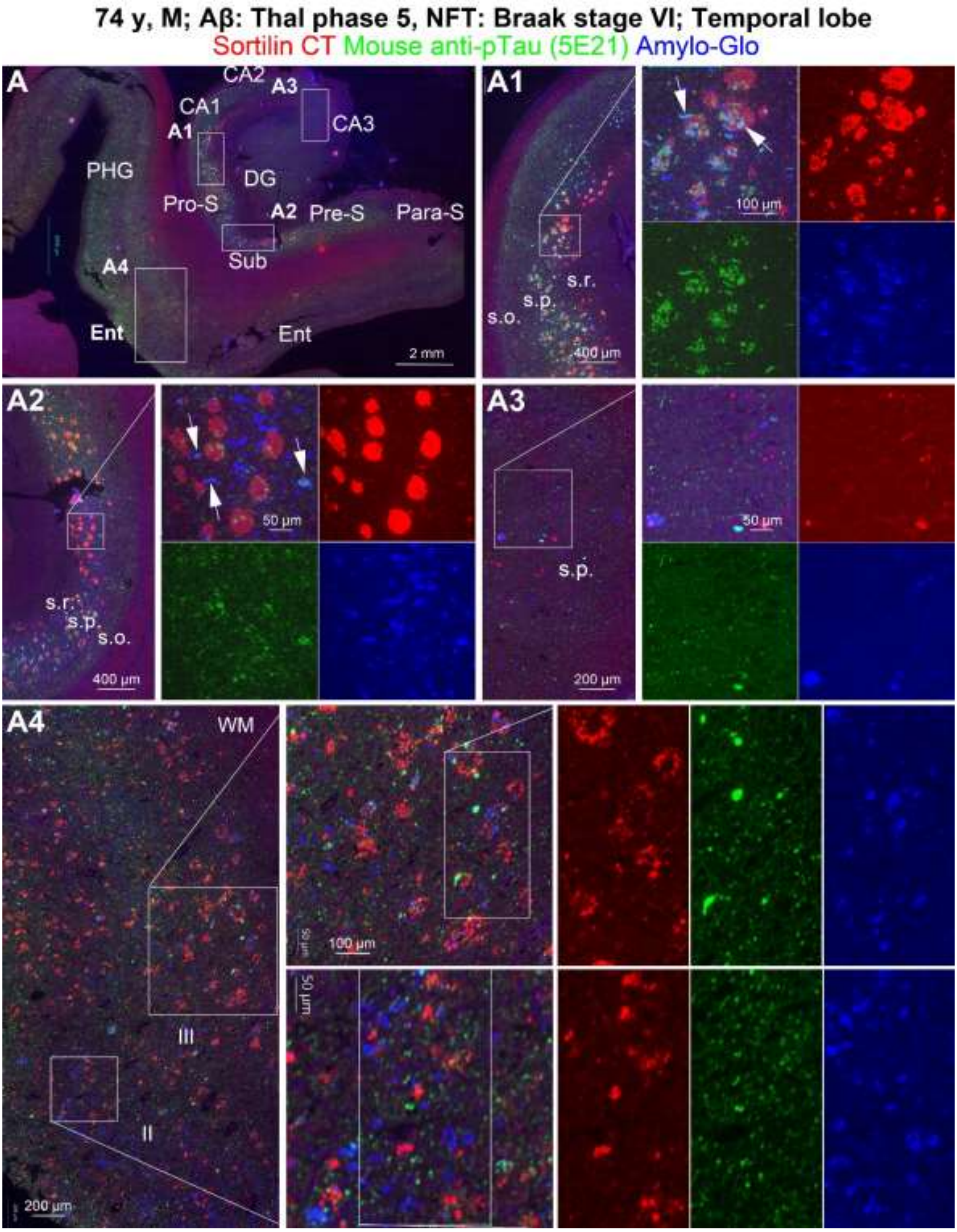


**Supplemental Figure 12: Tripe fluorescent characterization of sorfra plaque colocalization with tangle labeling with an additional mouse pTau antibody and Amylo-Glo counterstain.** Boxed areas in the low power image (**A**) are enlarged as indicated. Sorfra plaques coexist with neuritic profiles labeled for pTau and/or Amylo-Glo in the CA1 and subicular areas and in the entorhinal cortex (**A1, A2, A3, A4**). Many somal and dendritic profiles are only labeled by Amylo-Glo, representing ghost tangles. In the entorhinal cortex, plaques show a differential labeling of sorfra and Amylo-Glo (**A4**). In CA3, few sorfra plaques are present, with a low amount pTau/Amylo-Glo labeled somal and neuritic profiles (**A3**).

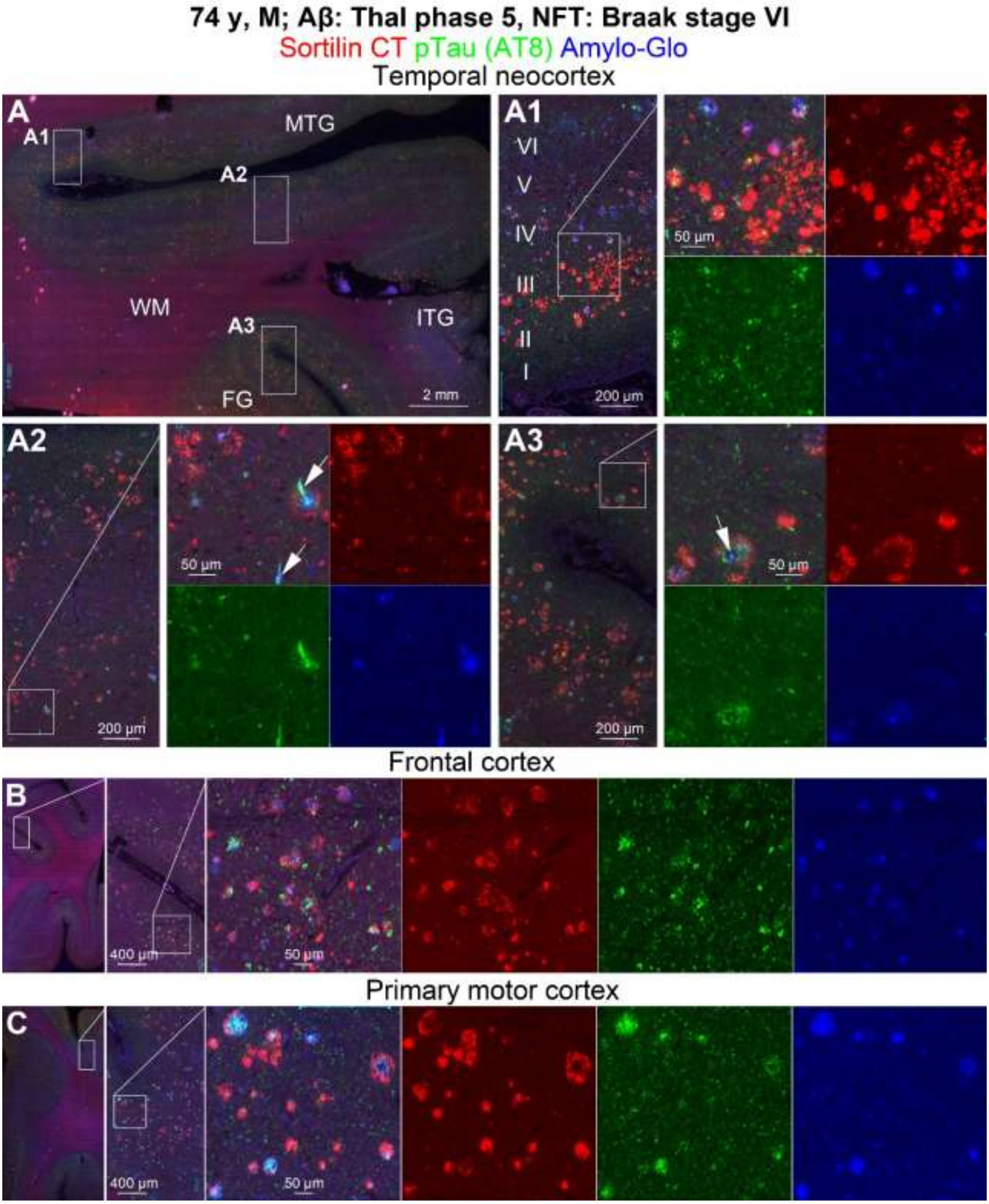


**Supplemental Figure 13: Tripe fluorescent characterization of sorfra plaque colocalization with tangle labeling with the AT8 pTau antibody and Amylo-Glo counterstain in cerebral neocortex.** Shown are images from the temporal (**A, A1, A2, A3**), frontal (**B**) and primary motor (**C**) cortices, with enlarged views arranged as indicated. Sorfra plaques are generally colocalized with neuritic elements labeled by AT8, Amylo-Glo or both (**A2, A3**, pointed by arrows). Some plaques show bright Amylo-Glo stain likely related to Aβ deposition in addition to tangle-bearing neurites (**C** and enlarged views).

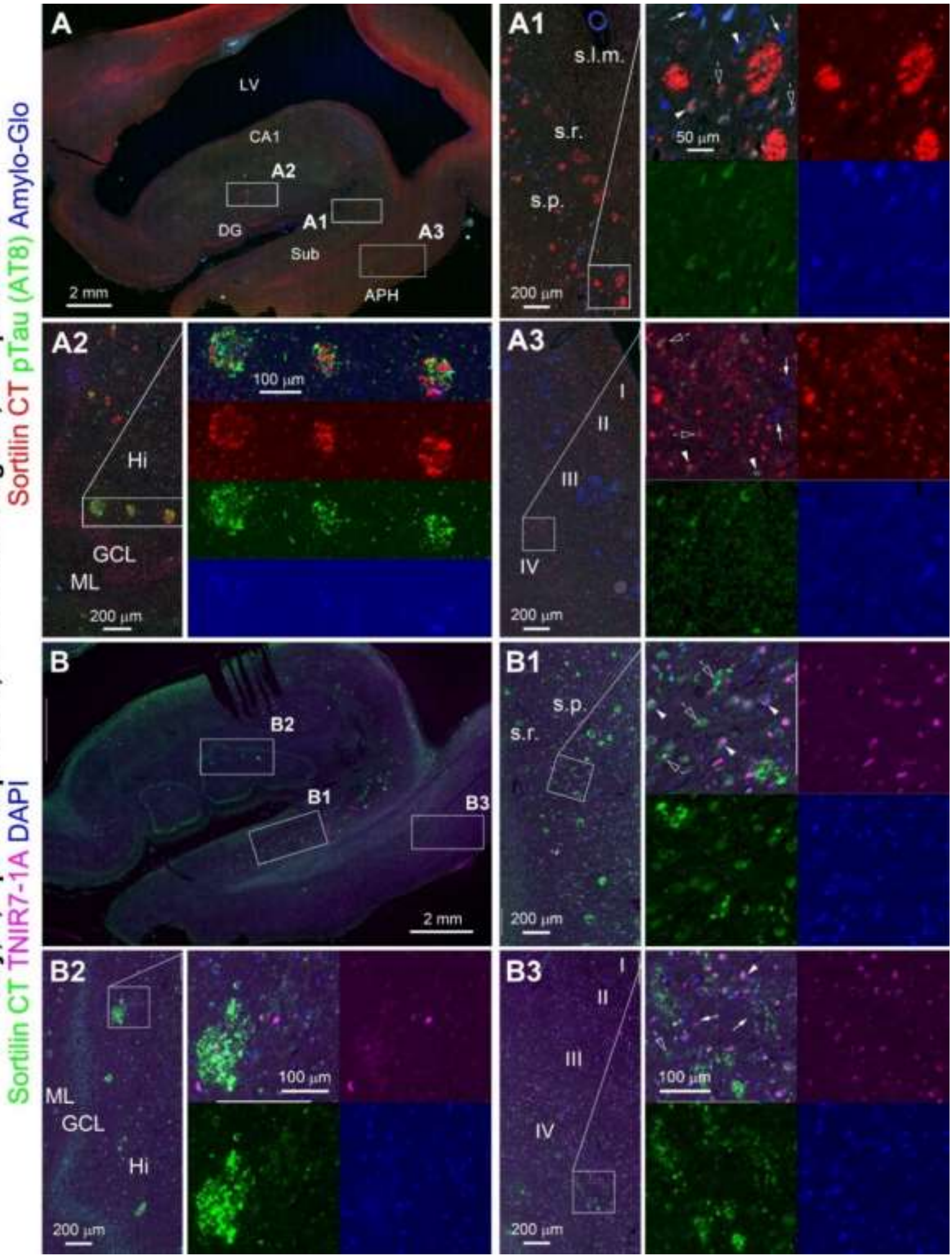


**Supplemental Figure 14: Comparative assessment of sorfra plaque colocalization with tangle labeling by the AT8 pTau antibody and a specific tangle tracer TNIR7-1A.** Massed sorfra plaques are associated with neuritic elements labeled by pTau/Amylo-Glo (**A, A1, A2**). Lightly stained sorfra plaques in the entorhinal cortex are in mix with neuritic threads, with mature (arrowheads) and ghost (arrows) tangles present nearby (**A3**). Sorfra plaques frequently colocalized with TNIR7-1A labeled neurites (**B, B1, B2, B3**). TNIR7-1A labeling also occurs inside neurons co-expressing sortilin (arrowheads), some with intracellular inclusion (open arrows) (**B1, B3**).

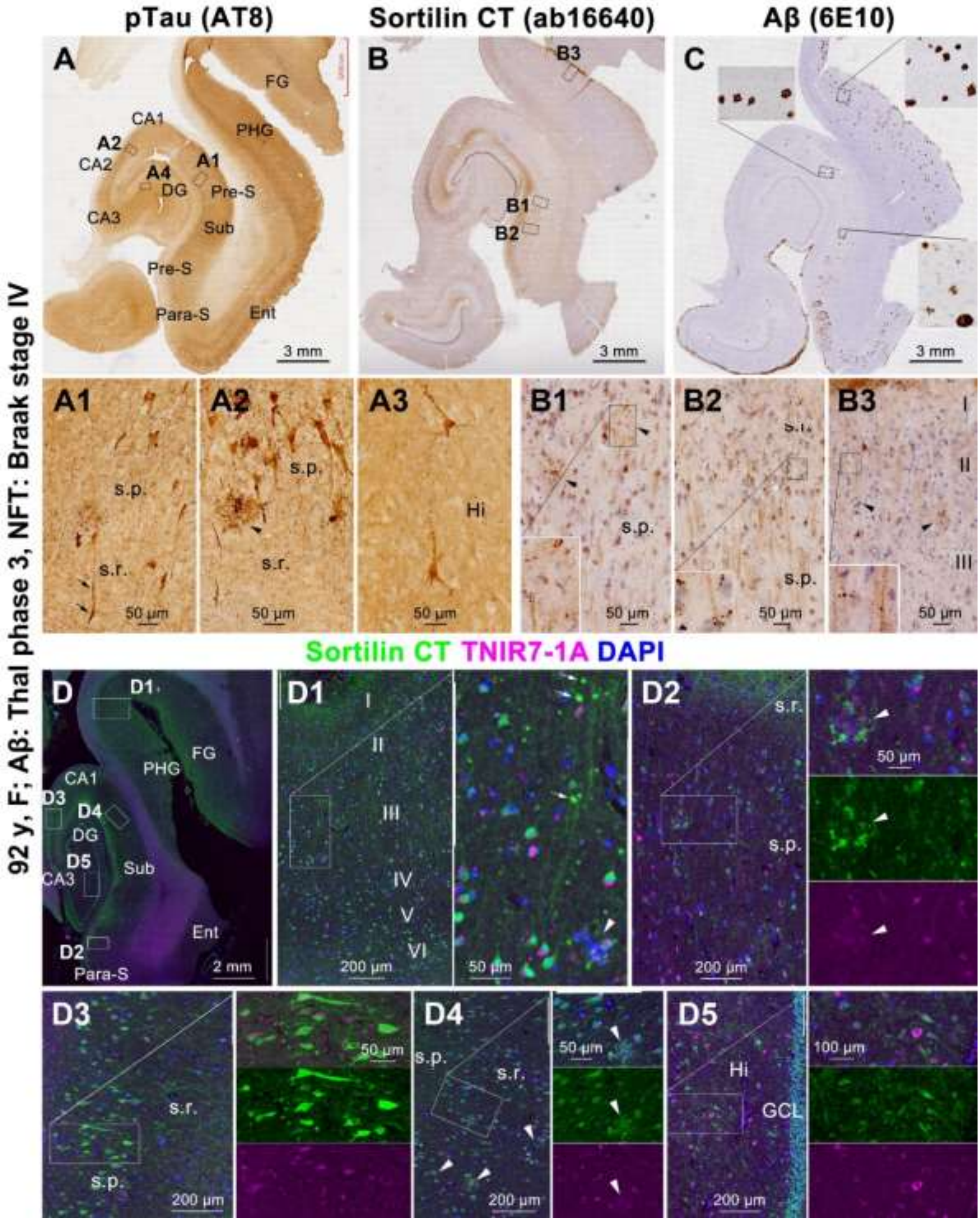


**Supplemental Figure 15: Assessment of early sorfra pathology relative to tangle lesions revealed by the TNIR7-1A tracer in temporal lobe sections with moderate AD pathologies**. pTau labeled neuritic clusters (**A, A1, A2**), sorfra plaques (**B, B1**) and Aβ plaques (**C,** inserts) occur in low numbers, whereas sortilin labeled neuronal somata and dendrites are present in all regions (**D, D1-D5**), some co-labeled by TNIR7-1A (enlarged views). Dot-like sortilin labeling is seen around the dendrites of pyramidal neurons (**B1-B3,** inserts; **D1,** arrows). Sorfra plaques (arrowhead) are associated with TNIR7-1A-labeled neurites (B3, D1, D2, D4). Intracellular inclusions (hollowed arrows) labeled by sortilin and TNIR7-1A can be found in some neuronal somata (**D1, D2, D3, D4, D5**, enlarged panels).

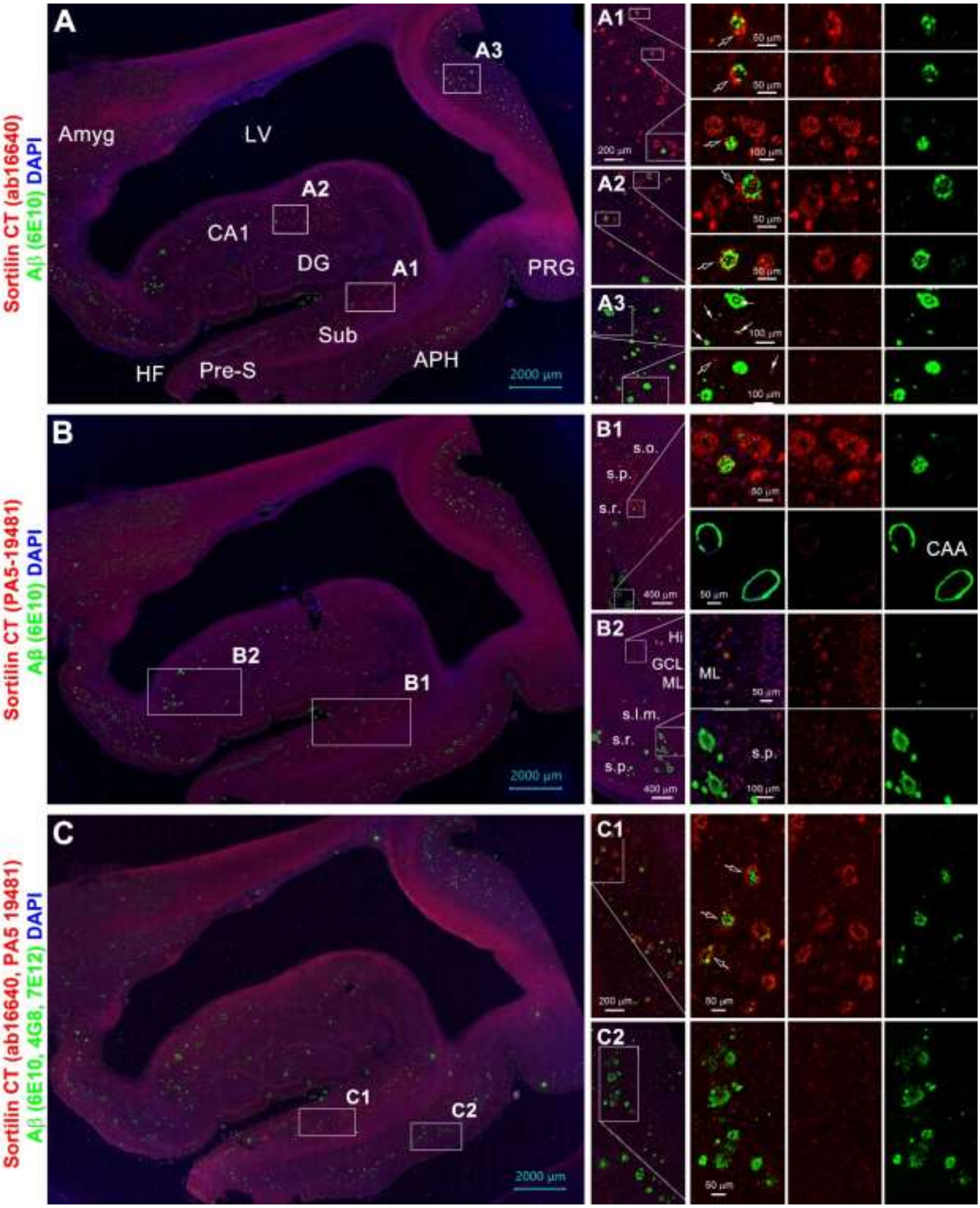


**Supplemental Figure 16: Double immunofluorescent characterization of sorfra and Aβ colocalization in temporal lobe sections using different antibody combination settings**. The antibody use settings are indicated on the left. Regardless of whether a pair (**A, B**) or a cocktail (**C**) of sortilin CT and Aβ antibodies are used, sorfra and Aβ colocalized and independent plaques are revealed (**A1, A2, A3, B1, B2, C1, C2**). The massed and cored compact-like Aβ plaques may not colocalize with sorfra (**A3**). Typical diffuse Aβ plaques and cerebral amyloid angiopathy (CAA) are not associated with sorfra deposition (**B1, C2**).

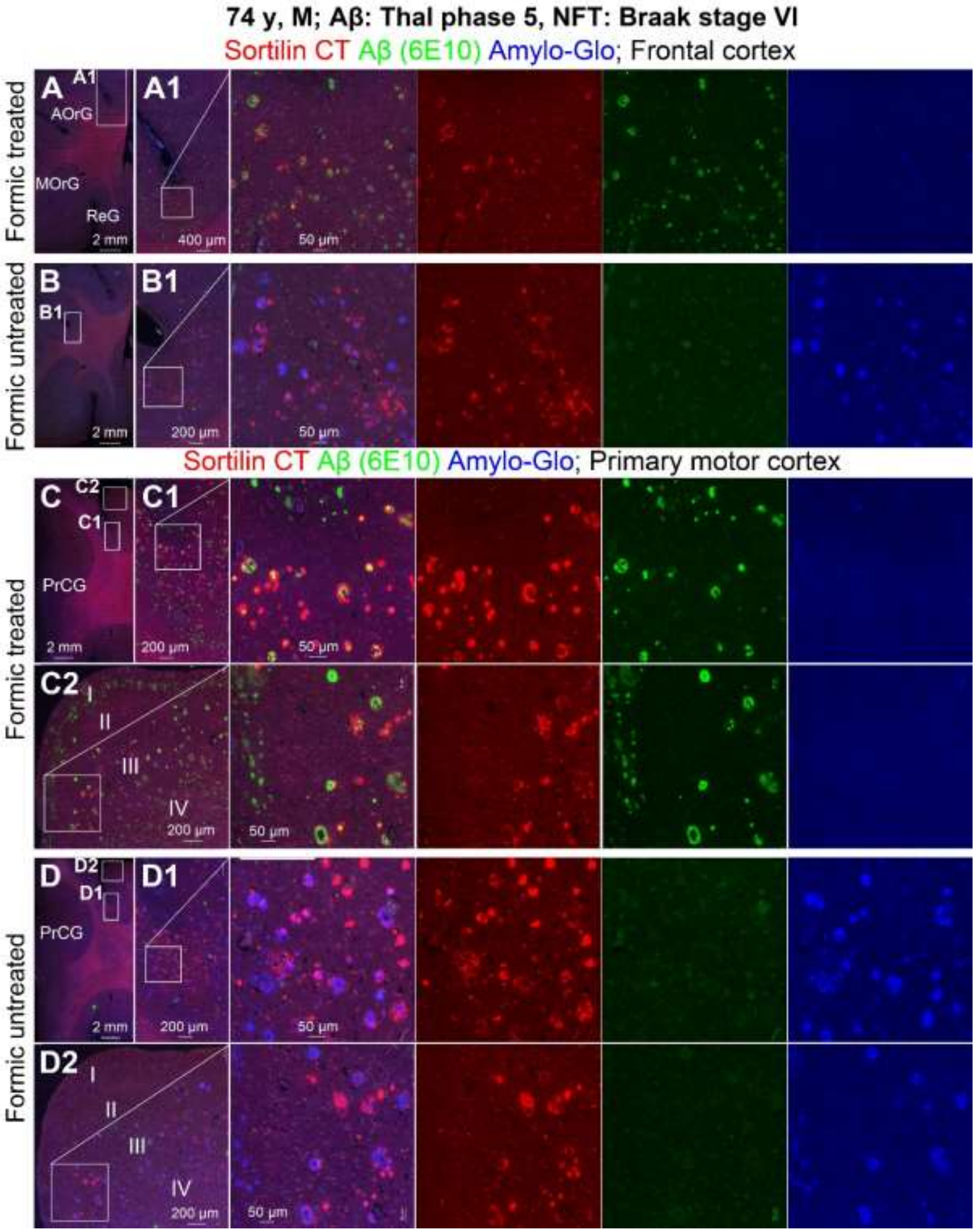


**Supplemental Figure 17: Double immunofluorescence of sorfra and Aβ with Amylo-Glo counterstain in neocortical regions with and without formic acid pretreatment of the paraffin sections**. A differential sorfra/Aβ colocalization was seen among plaque profiles in sections with formic acid treatment (A, A1, C, C1, C2). A differential sorfra/Amylo-Glo colocalization was seen among plaque profiles in sections without formic acid treatment (B, B1, D, D1, D2). In either condition, sorfra deposits often occur surrounding the Aβ/amyloid stain. Images are from the sulcal valley areas, with many plaques exhibited the sorfra immunolabeling only.

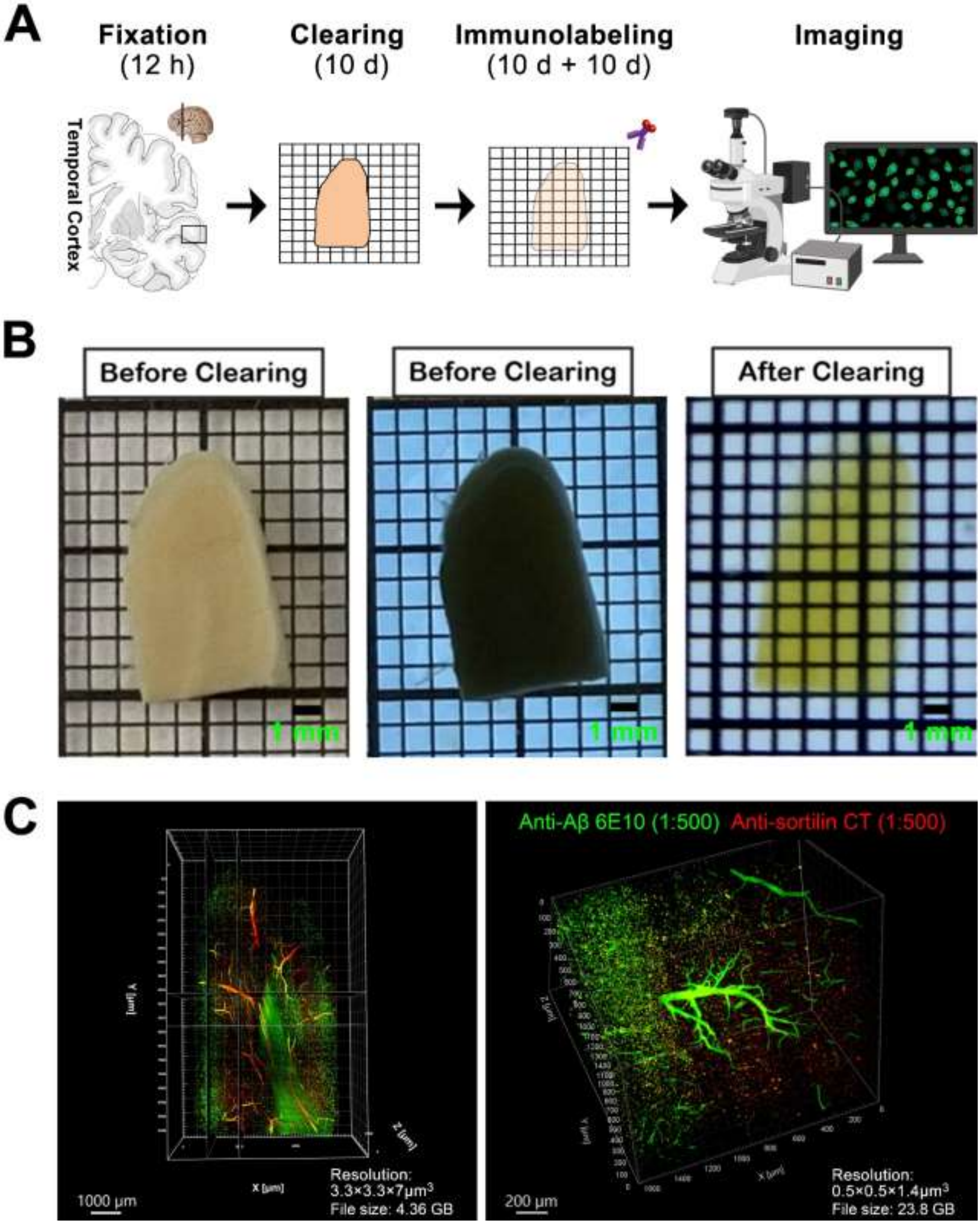


**Supplemental Figure 18: Protocol of cortical slice clearance, double immunofluorescence and imaging documentation**. Tissue information, experimental procedure, photo documentation and antibody use are as indicated (**A, B**). Panel (**C**) is the screenprints of the three-dimensional imaging video scanned at low and high resolutions as indicated, with a rendering of the vascular profiles in the tissue. The videos are provided as a separate supplemental file.

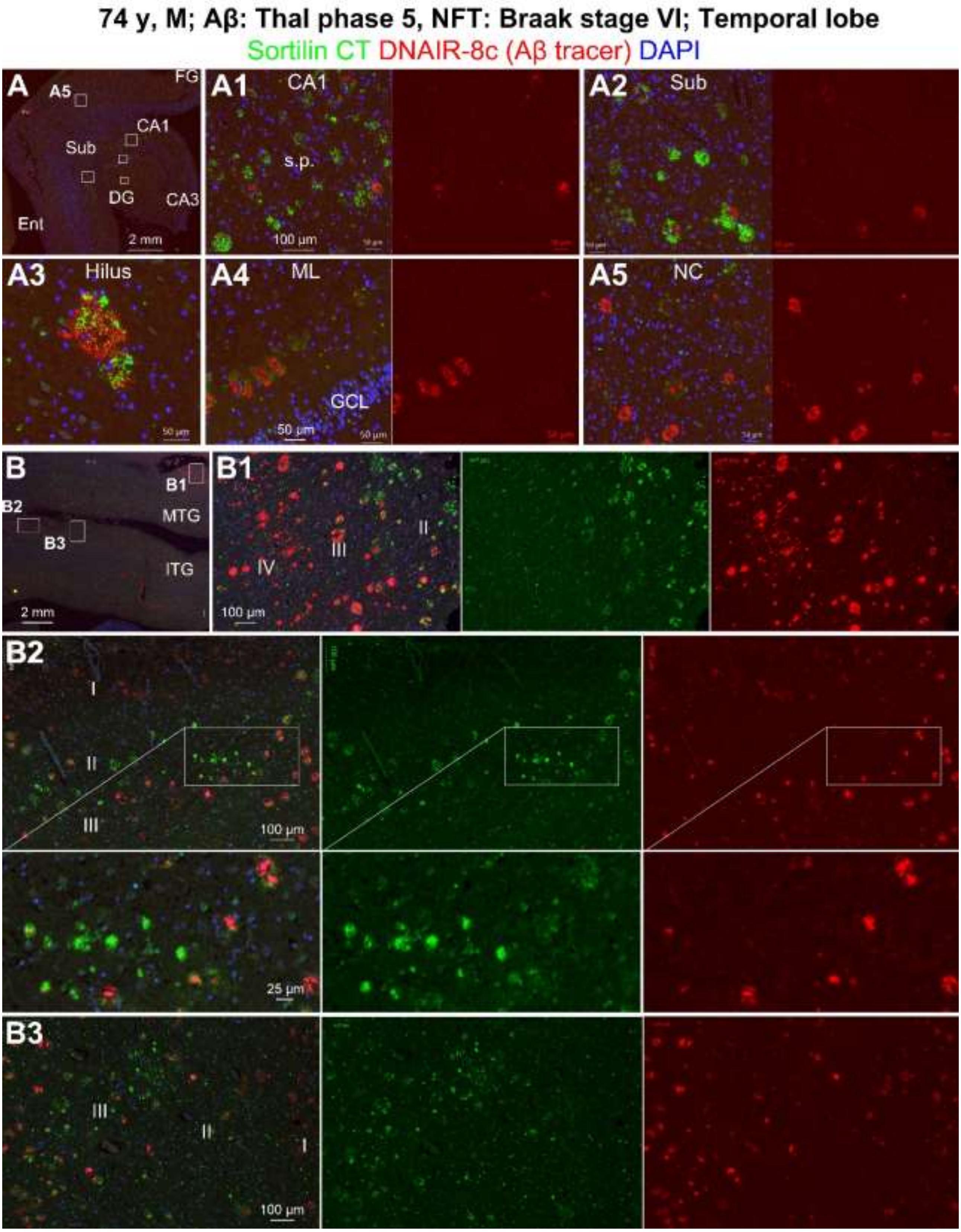


**Supplemental Figure 19: Characterization of differential sorfra and Aβ deposition as plaque profiles using a selective amyloid tracer, DNAIR-8c**. Mixed as well as independent plaques are present in various hippocampal subregions (**A, A1, A2, A3**) and cortical subregions (A5, B, B1, B2, B3) in a temporal lobe section from a case with end-stage AD pathology, as indicated.

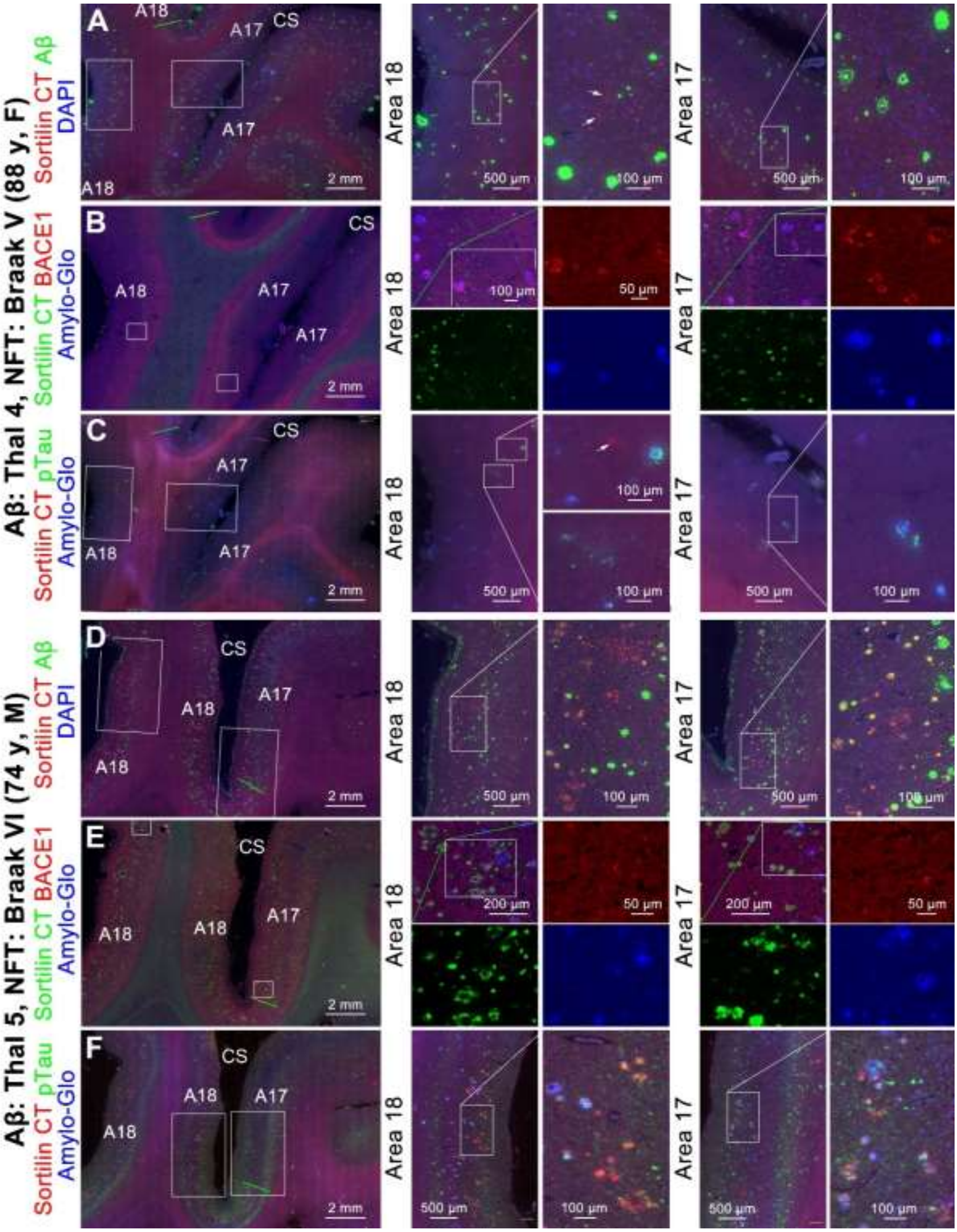


**Supplemental Figure 20: Parallel assessment on sorfra, Aβ, pTau and BACE1 immunofluorescent labeling in the primary and secondary visual cortical areas relative to Braak neurofibrillary tangle stages.** Aβ plaques and BACE1 labeled presynaptic dystrophic neurites occur in areas 17 and 18 at Braak stages V and VI (**A, B, D, E**). Sorfra and pTau labeling are occasionally seen at Braak stage V but frequently seen at Braak stage VI in areas 17 and 18, especially the former (**A, B, C, D, E, F**). Differential sorfra/Aβ colocalization (**D**) and frequent sorfra/pTau colocalization (**F**) in plaque profiles is seen at Braak stage VI.

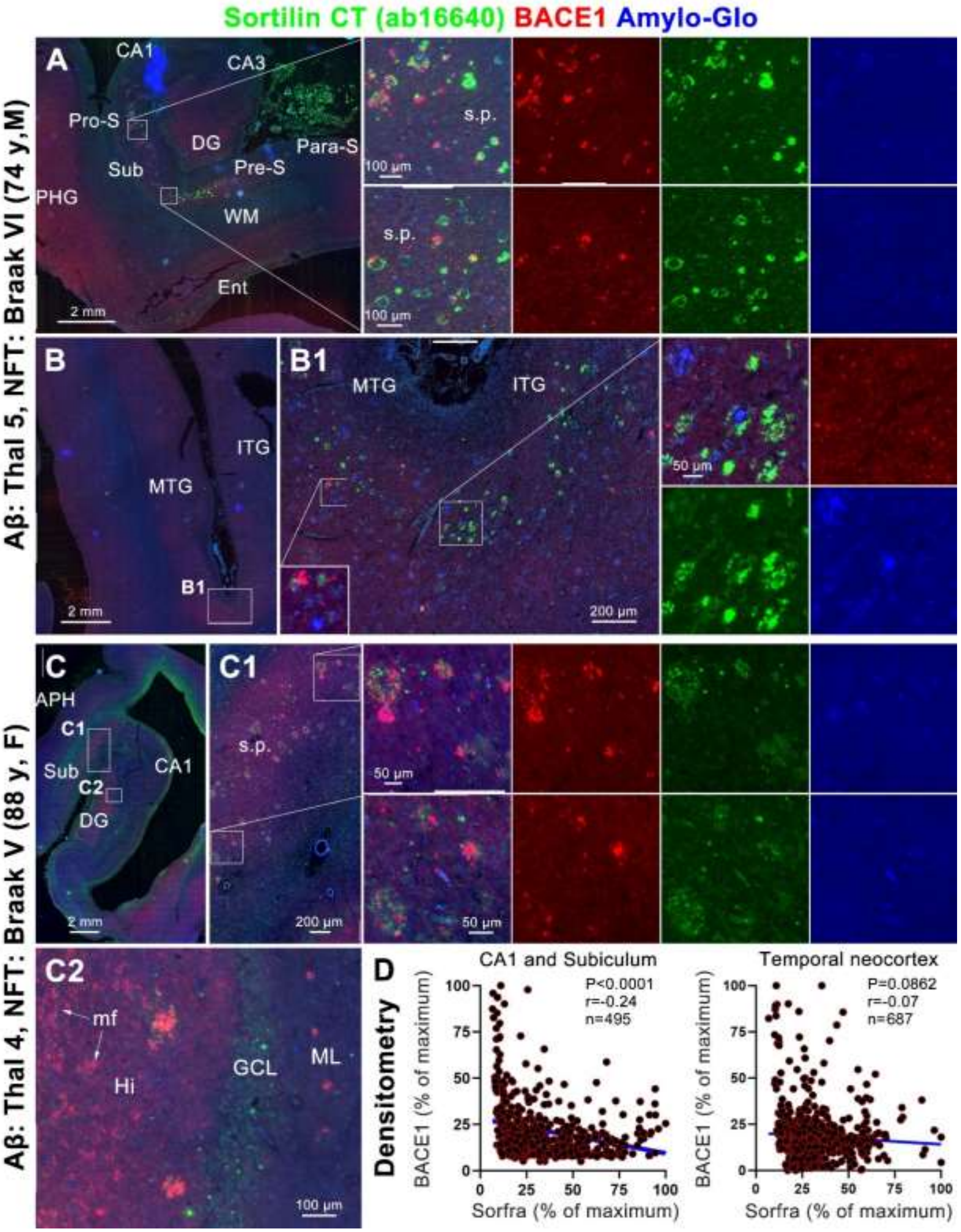


**Supplemental Figure 21: Microscopical and densitometric characterization on sorfra deposition relative to BACE1 labeled dystrophic neurite.** Image panels (**A, B, B1, C, C1, C2**) show a differential sorfra/BACE1 colocalization among the plaque profiles with Amylo-Glo stain in various hippocampal and cortical areas as indicated. BACE labeling is not seen in pure sorfra plaques. The dot graphs (D) show a negative correlation between sorfra and BACE1 labeling intensity among the colocalized plaque profiles quantified in subiculum/CA1 areas and in the temporal neocortex. mf: mossy fiber: mossy fiber.

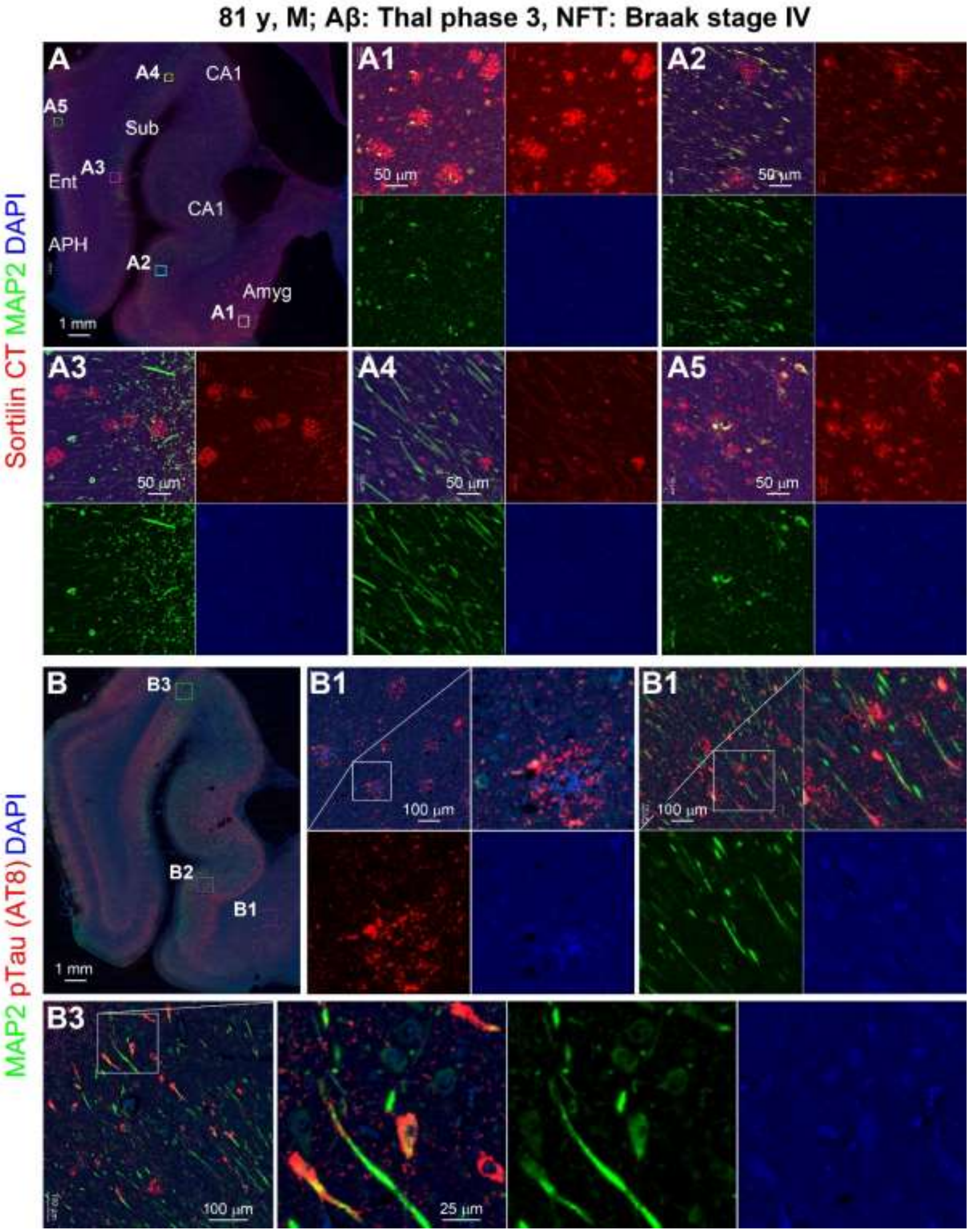


**Supplemental Figure 22: Association of sorfra and pTau pathology with reduced microtubule associated protein 2 (MAP2) labeling.** Sorfra plaques are associated with diminished MAP2 labeling regionally or locally, with sortilin/MAP2 co-labeled somatodendritic profiles existing in plaque-free areas and surrounding the plaques (**A, A1-5**). Colocalized neuritic remnants exist in some plaques (**A1, A5**). pTau labeled neuritic clusters generally lack MAP2 labeling (**B, B1, B2**). However, pTau/MAP1 labeling can coexist in the soma and dendrite in a complementary manner among some neurons (**B2, B3**).

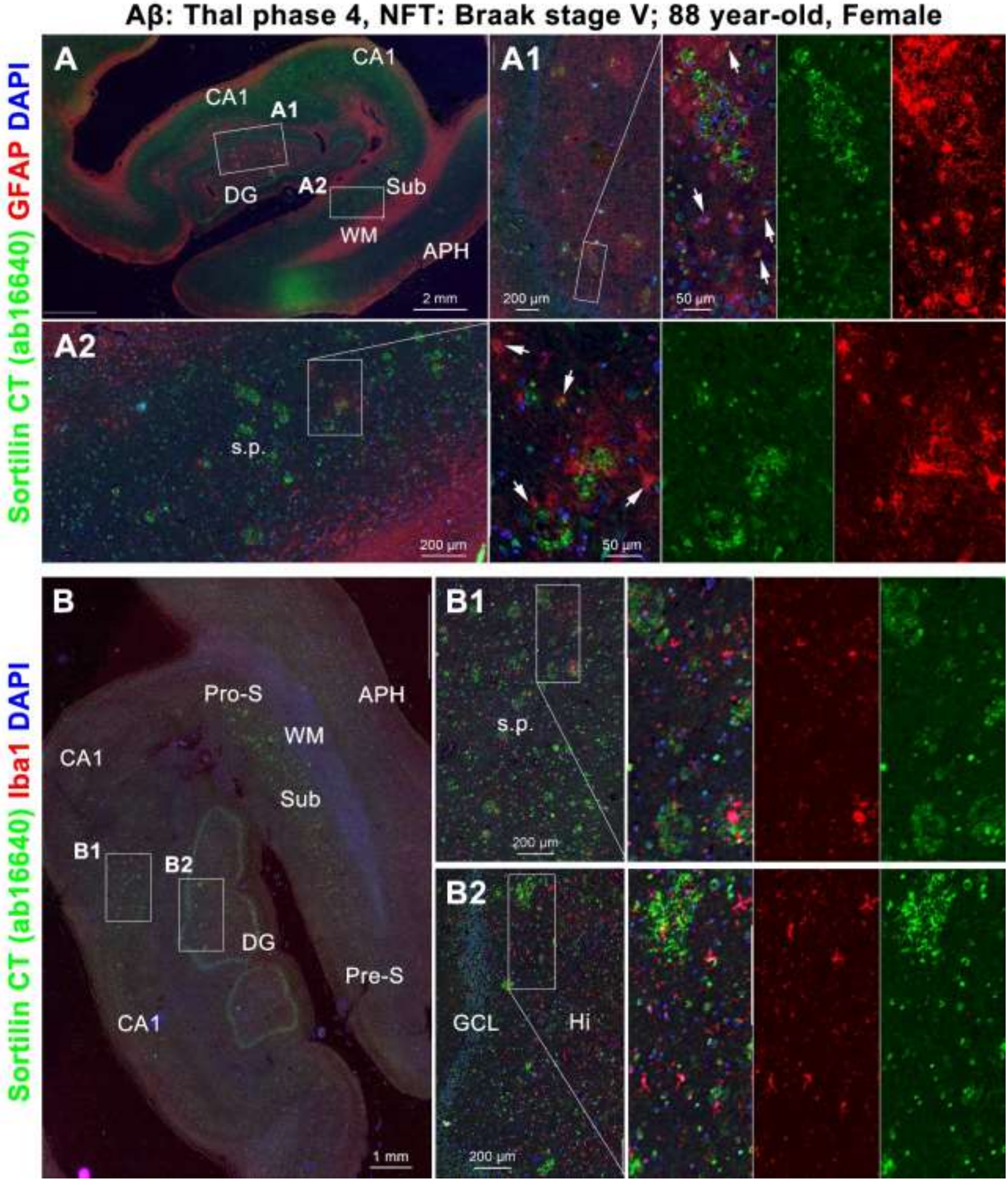


**Supplemental Figure 23: Double immunofluorescent characterization on potential sorfra phagocytosis by glial cells.** Shown are adjacent temporal lobe paraffin sections passing the hippocampal head from a brain with moderate AD pathology as indicated. Sorfra immunofluorescence is seen inside some astrocytes (pointed by arrows) labeled by a glial fibrillary acidic protein (GFAP) antibody (**A, A1, A2**) in the vicinity of the sorfra plaques. However, sorfra labeling is not seen in microglial cells immunolabeled by the Iba-1 antibody (**B, B1, B2**).

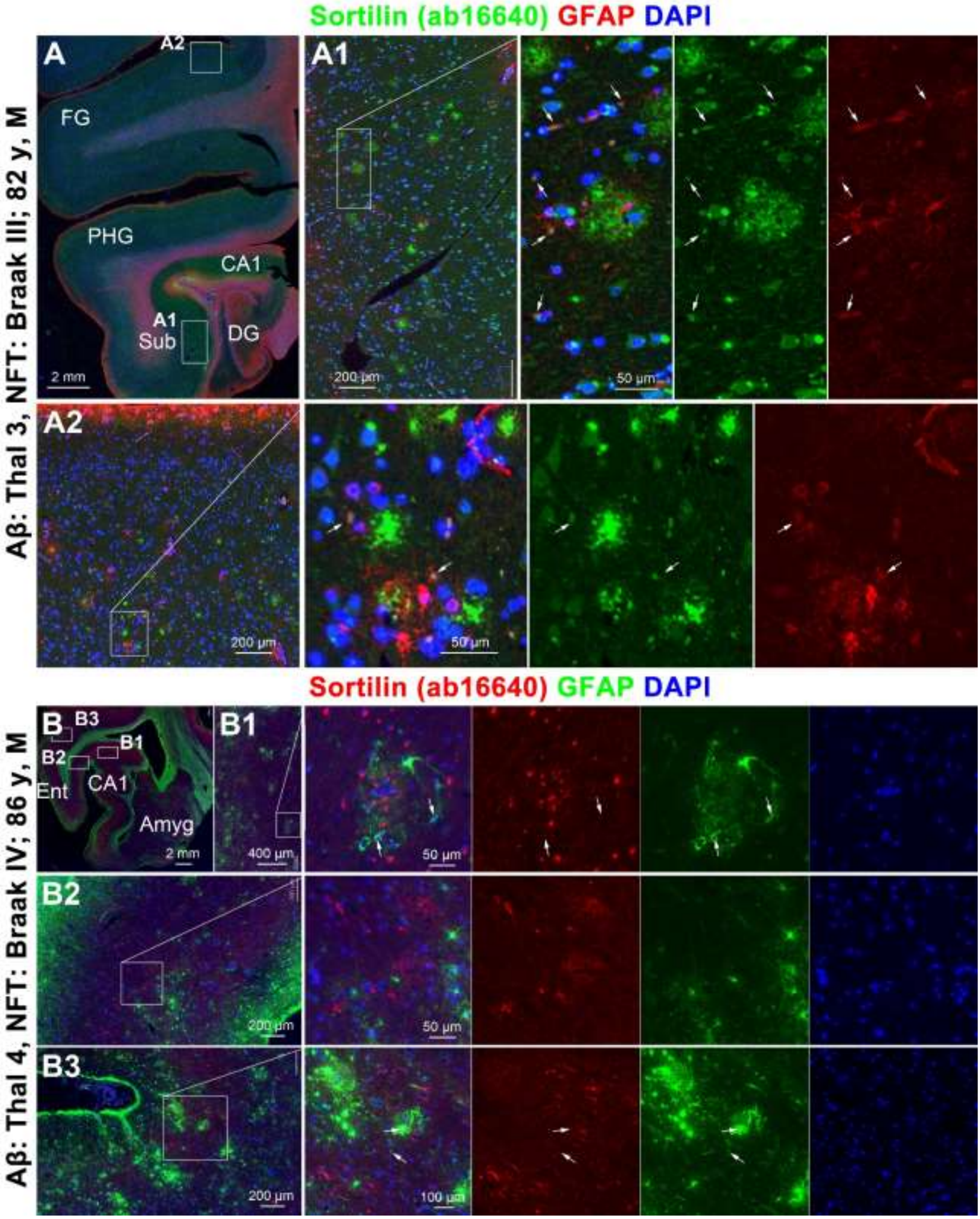


**Supplemental Figure 24: Additional examples showing sorfra labeling inside astrocytes.** Shown are temporal lobe paraffin sections from two brains with AD pathologies as indicated (**A, B**). There is no consistent association between the sorfra plaques and GFAP labeled astrocytes in anatomical location or in quantity. At the section cutting level, small dot-like sorfra labeling (pointed by arrows) could be identified inside some astrocytes around the sorfra plaques (**A1, A2, B1, B2, B3**).

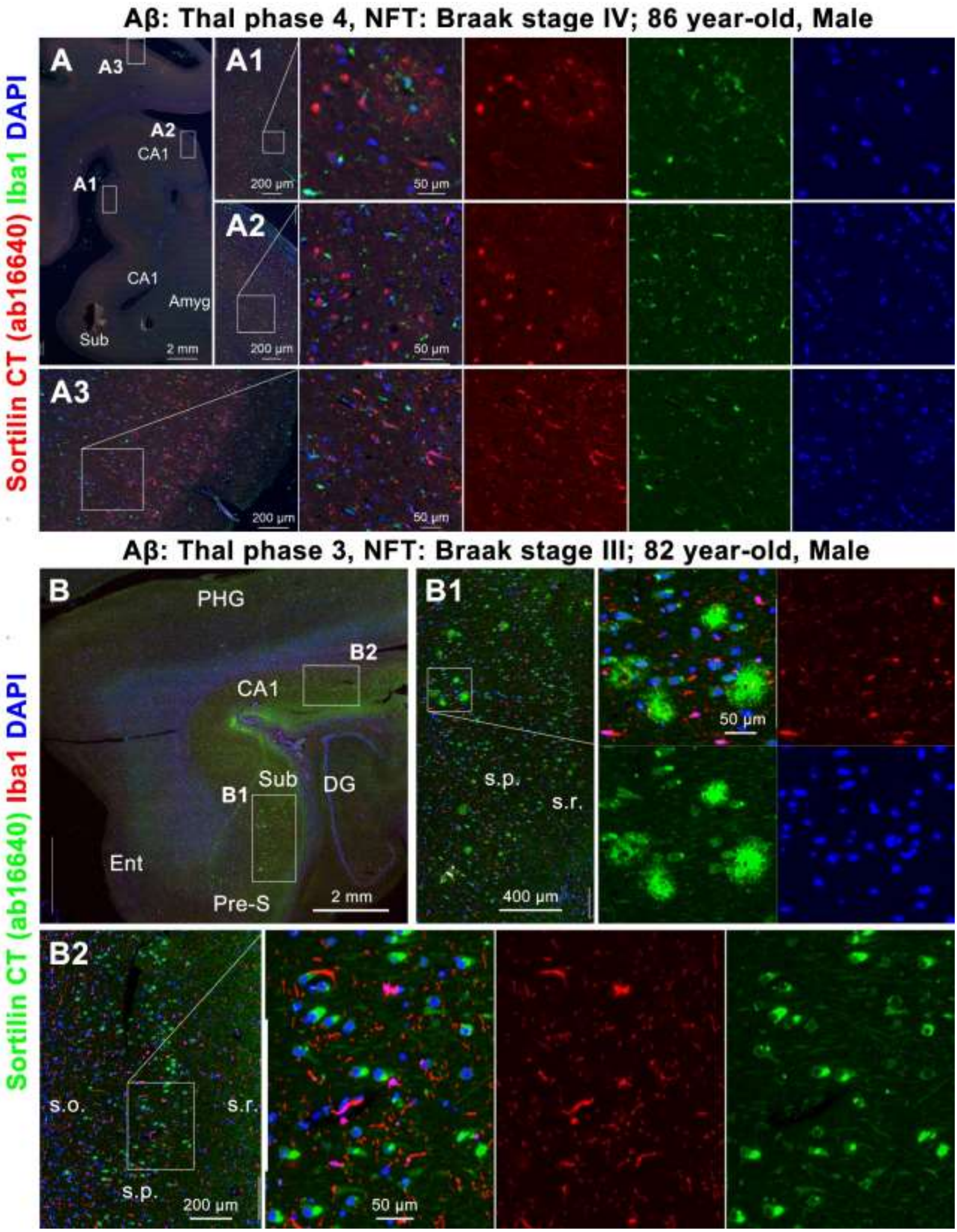


**Supplemental Figure 25: Additional examples showing the absence of sorfra colocalization with microglia.** Shown are temporal lobe paraffin sections from two brains with pathological scores as indicated (**A, B**). No colocalization of sorfra deposits in Iba1 labeled microglia cells is observed in the hippocampal formation or the cortical regions by close microscopic examination (A1, A2, B1, B2).
